# Spatial Metagene Discovery and Associated Molecular Pattern Characterization in Spatial Transcriptomics and Multi-Omics using SEPAR

**DOI:** 10.1101/2025.04.05.647412

**Authors:** Lei Zhang, Ying Zhu, Shuqin Zhang

## Abstract

Spatially resolved transcriptomics (SRT) has transformed biomedical research by enabling gene expression profiling at near- or sub-cellular resolution while preserving spatial context. However, interpreting SRT data to understand cellular and gene organization remains challenging. Current methods focus on identifying global spatial domains across all genes, but often miss localized structures driven by specific gene subsets. Here, we introduce SEPAR, an unsupervised computational framework designed to analyze SRT data using a limited number of spatial metagenes and their expression patterns. SEPAR integrates gene activity and spatial neighborhood relationships to summarize gene expression per spot/- cell, decomposing data into contributions from spatial metagenes. It supports downstream analyses, including metagene expression pattern-specific gene identification, spatially variable gene (SVG) detection, spatial domain delineation, and gene expression refinement. Applied to diverse datasets, SEPAR demonstrates high efficacy. Gene sets linked to spatial metagene patterns are enriched with meaningful cell types and gene ontologies. SVGs are detected with higher accuracy, and gene refinement enhances biological signals, improving correlation analysis of functionally related genes. In spatial multi-omics data, SEPAR excels at revealing strong correlations between co-localized molecules in spatial CITE-seq data and identifies coordinated gene-peak relationships in MISAR-seq data, offering insights into spatial molecular interactions.

## 1 Introduction

Spatially resolved transcriptomics (SRT) technologies, ranging from spot-based capture techniques to high-resolution imaging methods such as 10 × Visium [1], Slide-seq [2, 3] and MERFISH [4], have revolutionized biomedical research by providing gene expression levels at near- or sub-cellular resolution while preserving their precise spatial localizations. This advancement enables researchers to investigate the spatial organization of various cell types, explore the cellular functions and cell-cell interactions, as well as examine the spatially distinct gene expression patterns [5–9]. Such insights contribute to a more comprehensive understanding of the complex biological systems and the tissue architecture. Leveraging the success of SRT technologies, spatial molecular profiling has evolved to simultaneously capture multiple molecular modalities recently. For example, spatial CITE-seq [10] combines transcriptome profiling with protein abundance measurement, while MISAR-seq [11] facilitates concurrent profiling of gene expression and chromatin accessibility (ATAC) with spatial resolution. These multi-omics measurements facilitate studies of the interplay between transcriptional regulation, protein expression, and chromatin states within their native tissue context, thereby offering deeper insights into cellular function and tissue organization.

Various methods have been developed to identify the spatial domains with similar gene expressions for SRT data, such as BayesSpace [12], STAGATE [13], BASS [14], and so on. While these methods excel in capturing global tissue architectures, they have serious limitations in resolving fine-grained spatial structures inherent in the data. A key issue is their focus on dominant expression patterns shared across all genes, overlooking localized patterns driven by specific gene subsets that may define functionally specialized niches or rare cell states [15]. To address these limitations, we adopt the concept of metagenes, which has been proven successful in traditional transcriptomics analysis [16, 17]. Metagenes, as weighted linear combinations of functionally related genes, represent coordinated transcriptional programs that can recover underlying biological processes or significant phenotypes [17]. Gene sets exhibiting similar spatial expression patterns can be systematically identified through their alignment with metagene expression patterns. This process can be achieved using matrix factorization methods, particularly Non-negative Matrix Factorization (NMF), which has been proven effective in extracting interpretable biological features [18]. Applications of NMF to single-cell RNA sequencing data [19–21] have also demonstrated its capability in revealing coordinated gene expression programs. However, these methods do not incorporate spatial information, limiting their utility for SRT data analysis. Recent advancements like GraphPCA [22] have attempted to incorporate spatial information into dimensionality reduction. Still, the linear nature of principal component analysis (PCA) and lack of non-negativity constraints make it challenging to obtain interpretable spatial patterns of gene groups. Similarly, probabilistic modeling approaches such as STAMP [23] employ deep generative models combined with simplified graph convolution networks to capture spatial dependencies in SRT data. However, the use of auto-encoding and black-box variational inference may impact model interpretability, and users are required to pre-specify the number of topics, which may require multiple iterations to optimize. Overall, neither approach can be naturally extended to multi-omics integration, a critical demand in spatial biology.

Identifying spatially variable genes (SVGs) that exhibit distinct spatial expression patterns is also crucial in SRT analysis. Different from the pattern-specific genes that show significant signals on particular metagene expression patterns, each SVG may cover one or more spatial patterns. By identifying SVGs, researchers can explore the relationship between spatial organization and molecular cell function, providing valuable insights for biologists and pathologists [24]. Furthermore, SVGs enhance analyses such as spatially-aware unsupervised clustering and trajectory inference, contributing to a better understanding of tissue morphology and inter-cellular communication [12, 25–28]. Current methods for identifying SVGs include SpatialDE [9], SPARK [29], SPARK-X [30], STAMarker [31], and so on. SPARK, SPARK-X, and SpatialDE utilize various statistical frameworks, such as Gaussian process regression and nonparametric approaches, to test the spatial variance and dependence of gene expression. However, these methods conduct hypothesis tests independently for each gene and fail to determine their corresponding spatial patterns. STAMarker, a model based on graph neural networks, relies on a pre-trained clustering model, making it difficult to identify the genes that do not contribute to clustering.

To address the above challenges in SRT data analysis, we developed a Spatial metagene Expression PAttern Recognition (SEPAR) method, a tool designed based on graph-regularized non-negative matrix factorization [32] to identify the distinct spatial expression patterns of metagenes. SEPAR integrates the gene expression and the spatial coordinates, capturing the complementary spatial expression patterns across genes while ensuring the computational efficiency. It offers several key advantages. First, SEPAR fully leverages the interpretability of NMF. It can extract meaningful spatial expression patterns of metagenes [17] and the pattern-specific genes, facilitating enrichment analysis of relevant genes and cells. Second, SEPAR can efficiently handle various downstream tasks such as SVG identification, unsupervised spatial clustering, and gene refinement. Third, SEPAR does not rely on prior assumptions on data distribution, allowing seamless extension to multimodal spatial data analysis. Last but not the least, SEPAR can be extended to multislice version, enabling simultaneous analysis of adjacent tissue sections. We evaluated SEPAR across various spatial molecular profiling datasets, including SRT data generated using 10× Genomics Visium [1], Slide-seqV2 [2, 3], Stereo-seq [33], MERFISH [4], and osmFISH [34], as well as spatial multi-omics data from spatial CITE-seq [10] and MISAR-seq [11]. Across all these platforms and modalities, SEPAR consistently demonstrated its efficiency in delivering robust analytical outcomes.

## 2 Results

### Overview of SEPAR

SEPAR is a novel framework based on graph-regularized non-negative matrix factorization [32], specifically designed to identify the underlying spatial metagenes and their expression patterns in SRT data (Fig. 1, **Methods**). The gene expression matrix is modeled as weighted combinations of metagenes, regularized by the terms that account for the spatial relationships of the spots and that enforce the metagene dissimilarity, ensuring clear and distinct patterns. The SEPAR framework is formulated as the following optimization problem:

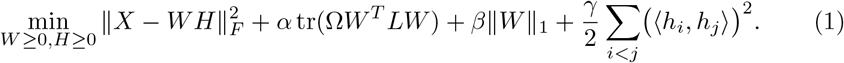

**Fig. 1:**
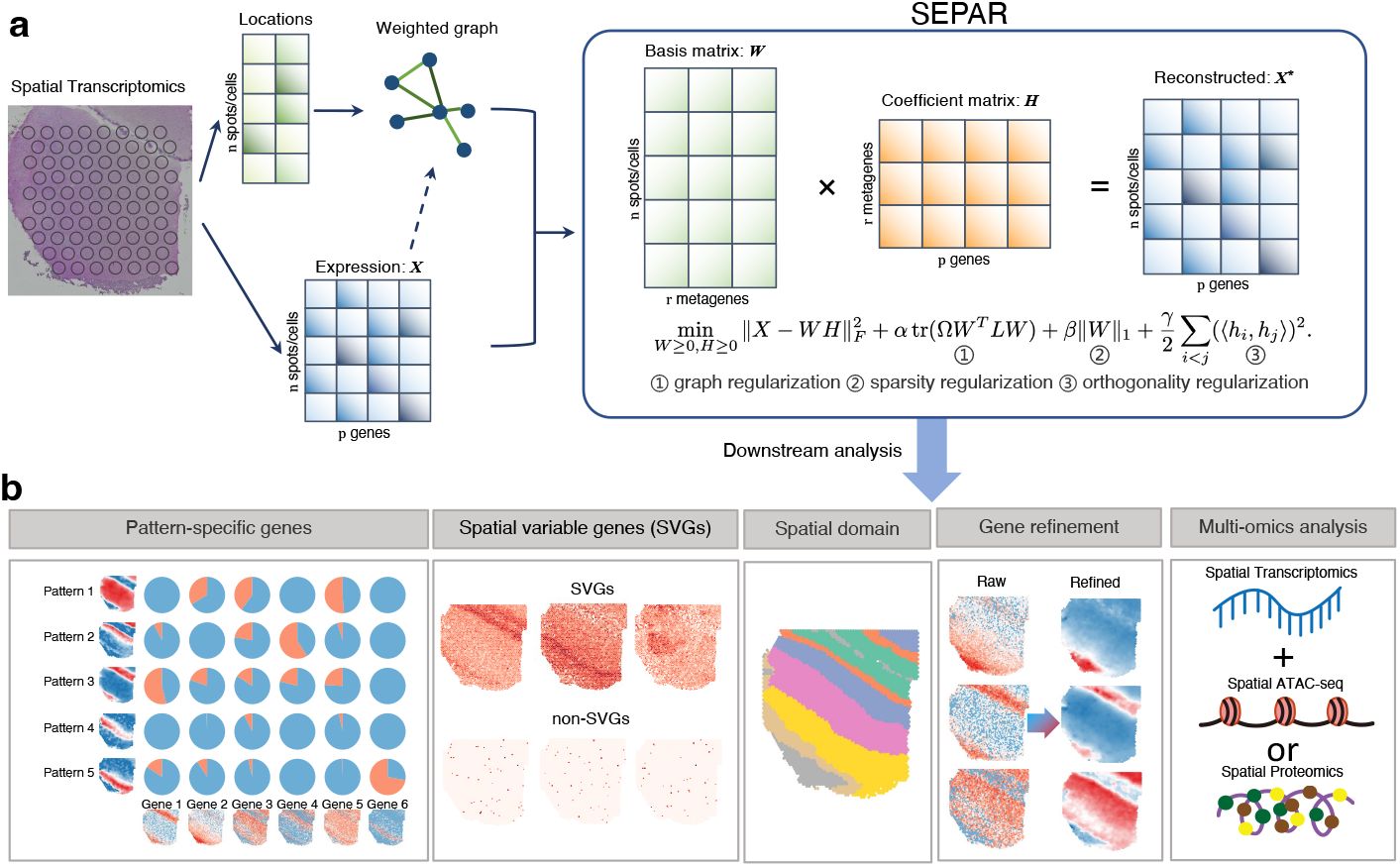
**(a)** SEPAR is a framework based on graph-regularized NMF designed to identify spatially aware metagene patterns using both spatial location and gene expression as input. Spatial location data is leveraged to construct weighted graph regularization capturing the spatial relationships of the spots or cells. Sparsity regularization and dissimilarity regularization are applied to ensures distinct spatial metagene patterns. **(b)** SEPAR supports efficient and robust downstream analyses of SRT data, including metagene expression pattern recognition, pattern-specific gene analysis, SVG identification, spatial domain delineation, gene expression denoising and spatial multi-omics data analysis.

Here, *X*_*n×p*_ denotes the gene expression matrix of *n* spots (or cells) and *p* genes. Matrix *W*_*n×r*_ represents the *r* underlying spatial metagene expression patterns in all the spots (or cells). And matrix *H* has size *r*× *p*, with each of the *r* rows defining a metagene. The spatial relationships of the spots/cells are incorporated using an adjacency graph in the second term. Ω represents the weights of the patterns derived from pattern significance scores, which prioritize patterns exhibiting strong spatial structure in the data. Sparsity regularization (*ℓ*_1_-norm) is introduced to mitigate noise effects, and the similarity term ∑_*i<j*_ (⟨ *h*_*i*_, *h*_*j*_⟩ )^2^ ensures distinct metagenes. This framework is also extended to multislice scenarios (**Supplementary Information**).

The results derived from SEPAR enable various downstream analyses, including spatial expression pattern recognition of metagenes, pattern-specific gene analysis, SVG identification, spatial domain delineation, gene expression denoising, and the analysis of spatial multi-omics data. The columns of matrix *W* represents the distinct spatial expression patterns of the metagenes. Pattern-specific genes, which are significantly expressed in particular spatial regions, are pinpointed by measuring the similarities between the given gene expression profiles and the underlying spatial meta-gene expression patterns. As SVGs can be expressed across multiple spatial metagene patterns, they are identified by evaluating the distance between each gene and combinations of the spatial patterns. Additionally, the decomposed patterns facilitate spatial domain identification by directly applying *k*-means to matrix *W* . The denoised gene expression matrix is reconstructed by combining *W* and *H*. By horizontally concatenating the spatial multi-omics data into a larger matrix *X*, SEPAR can be directly applied to infer the underlying structures and associated molecular signatures.

We comprehensively evaluated the performance of SEPAR across diverse spatial transcriptomics datasets and multiple scenarios. We first demonstrated SEPAR’s capabilities on the DLPFC dataset generated using 10× Visium [1] and the mouse olfactory bulb dataset generated by Stereo-seq [33], where it effectively identified biologically meaningful spatial metagene patterns and SVGs. For image-based spatial transcriptomics, we validated the performance of SEPAR using datasets generated by MERFISH [35] and osmFISH [34], focusing on spatial domain identification and gene expression refinement. Furthermore, we extended SEPAR’s application to multiomics scenarios, demonstrating its versatility with the datasets generated by spatial CITE-seq [10] and MISAR-seq [11], highlighting its robustness in managing complex spatial molecular data. In addition, SEPAR can be also extended to handle multi-slice SRT data, which is demonstrated with the DLPFC dataset [1]. These extensive experiments illustrate that SEPAR not only achieves superior or competitive performance compared to existing methods but also offers computational efficiency and broad applicability across different spatial molecular profiling platforms.

### 2.1 SEPAR reveals layer-specific patterns and robust SVGs in human dorsolateral prefrontal cortex

The dorsolateral prefrontal cortex (DLPFC) dataset [1] was generated using the 10× Visium platform. It consists of a total of 12 slices with manual annotations of the DLPFC layers and white matter (WM) (Fig. 2a). Each slice contains over 3000 spots, with over 20000 genes measured. This dataset is widely utilized for evaluating spatial domain identification techniques, and encompasses many biologically significant SVGs linked to the structural layers of the cortex [1].

**Fig. 2:**
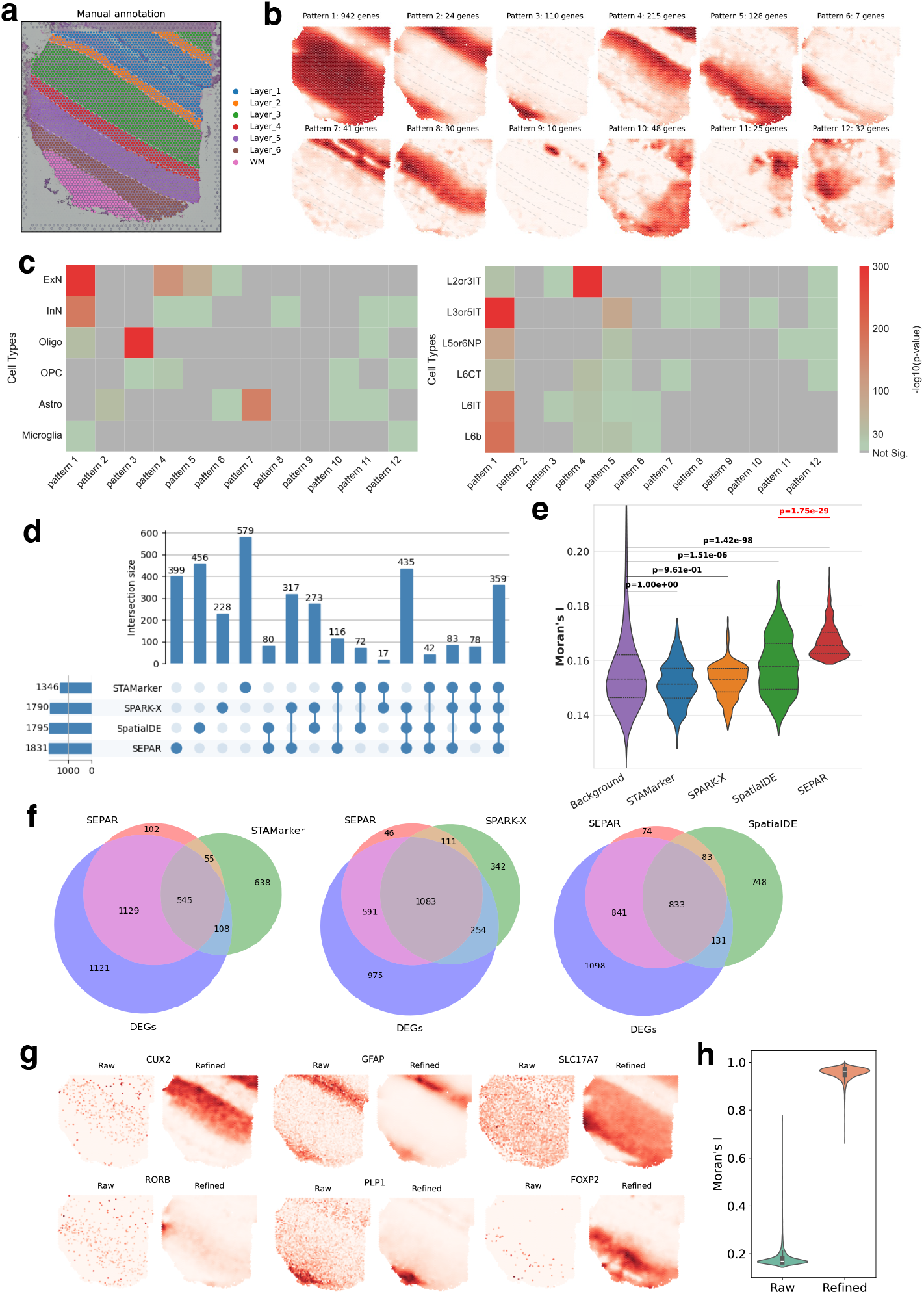
Analysis of section 151507 of DLPFC dataset generated by 10× Visium using SEPAR. (**a**) Manual annotations of section 151507 from original research [1]. (**b**) 12 spatial metagene patterns recognized by SEPAR with number of pattern-specific genes. (**c**) Cell-type enrichment analysis on the gene sets corresponding to each pattern by CellGO [36], using identifier with cell types including ExN, InN, Oligo, OPC, Astro and Microglia (left) and with ExN cell including L2/3IT, L3/5IT, L5/6NP, L6CT, L6IT, and L6b (right). (**d**) UpSet plot for SVGs identified by STAMarker, SPARK-X, SpatialDE and SEPAR. (**e**) Comparison of Moran’s I values for uniquely identified SVGs by each method. Background genes are all genes excluding the common SVGs identified by the four methods. One-sided Mann-Whitney U test is used to calculate the p-values. (**f** ) Venn plot comparing the SVGs from four methods and the DEGs selected by DEseq2. SVGs identified by SEPAR show significantly greater overlap with DEGs than those detected by STAMarker, SPARK-X, or SpatialDE. (**g**) An illustration showcasing the effects of SEPAR refinement on 6 marker genes. (**h**) Violin plots comparing the distribution of Moran’s I values for genes before and after refinement.

SEPAR can successfully identify the significant spatial metagene expression patterns and determine the specific genes associated with each pattern. Fig. 2b highlights the 12 key patterns selected according to the pattern significance score (PSS), along with the number of pattern-specific genes. The smallest number of associated genes is 7 and the largest is 942. We examined how each pattern maps to the manually annotated cortex layers (Fig. S1). The violin plots illustrate the distribution of 12 spatial meta-gene expression patterns across manually annotated cortical layers (L1-L6) and white matter (WM), revealing the relationship between computationally extracted expression patterns and anatomically defined laminar organization. Patterns 1-8 exhibit a strong correspondence with the structural cortical layers. For instance, Pattern 2 correlates with layers L1 and WM, while Pattern 3 corresponds predominantly with WM. Notably, Patterns 10-12 reveal novel spatial organizations that do not conform to the classical laminar structure of DLPFC. Subsequent cell-type enrichment and gene ontology analyses confirm the biological significance of these non-canonical patterns (Fig. 2c, Fig. S2, Fig. S3), which reveal their roles in extracellular organization, lipid metabolism, and membrane specialization, respectively. This demonstrates SEPAR’s ability in detecting meaningful metagene patterns beyond conventional layer-based analysis.

Cell-type enrichment analysis of the metagene expression pattern-specific genes shows distinct cellular compositions across spatial domains. Using CellGO [36] with three established cell-type identifiers [37], we examined the cellular preferences of each pattern. Analysis with major brain cell populations (ExN, InN, Oligo, OPC, Astro, and Microglia) and detailed excitatory neuron subtypes (L2/3IT, L3/5IT, L5/6NP, L6CT, L6IT, and L6b) in the prefrontal cortex revealed cell-type-specific enrichment results (Fig. 2c). Patterns 1 and 4 are significantly associated with excitatory neurons (ExN), with p-values of 1.65e-294 and 5.07e-130, respectively. Further analysis revealed layer-specific preferences, where Pattern 1 showed stronger enrichment for L3/5IT cells (p-value = 0), while Pattern 4 exhibited preferential enrichment for L2/3IT cells (p-value = 0), consistent with the spatial distribution of these patterns. Pattern 3 shows strong enrichment for oligodendrocytes (p-value = 0) and significant association with L6IT excitatory neurons (p-value = 7.56e-19), suggesting important neuron-glia interactions specific to deeper cortical layers. Additional analysis of inhibitory neuron subtypes (LAMP5, PVALB, SST, VIP, and SNCG) is presented in Fig. S2.

Gene-ontology enrichment analysis of pattern-specific genes reveals distinct functional characteristics of each identified patterns (Fig. S3). Using g:Profiler [38], we found Pattern 10 shows significant enrichment in terms related to extracellular space (p-value = 2.15e-07) and anatomical structure morphogenesis (p-value = 8.27e-05). This gene set and the corresponding structure likely involve components of the extra-cellular matrix, growth factors, and signaling molecules, all of which are ubiquitously distributed and essential for maintaining the structure and function of the cortex. Pattern 11 reveals significant enrichment in lipid metabolic processes (p-value = 2.49e-13). Additional enrichment in cellular lipid metabolic processes, regulation of lipid metabolism, and lipid catabolism (all p-values *<*7.30e-09) further underscores the critical role of lipid metabolism in this brain region. These findings reveal a distinct region predominantly in layer L1 of DLPFC with enhanced lipid catabolic activity, suggesting the presence of specialized cellular populations with elevated lipid metabolic processes, which may contribute to local neuronal function and energy homeostasis [39]. We note that Pattern 10 and 11 were not identified by existing methods.

SPEAR can identify the SVGs with high accuracy. We compared with the updated SVG identification methods: STAMarker [31], SPARK-X [30], and SpatialDE [9]. The UpSet plot in Fig. 2d visualizes that 359 SVGs were consistently identified across all four methods, which underscores significant concordance. Notably, the greatest three-way overlap was between SEPAR, SpatialDE, and SPARK-X, with 435 common SVGs, indicating substantial agreement for specific SVG types. To further evaluate the performance of these methods, we calculated the spatial autocorrelation of uniquely identified SVGs using Moran’s I [40]. As shown in Fig. 2e, SVGs identified by SEPAR and SpatialDE exhibited significantly higher Moran’s I values (p-value = 1.42e-98 and p-value = 1.51e-06, respectively; one-sided Mann-Whitney U test) compared to background genes, with SVGs identified by SEPAR showing significantly higher Moran’s I values than those obtained using SpatialDE (p-value 1.75e-29). The SVGs uniquely identified by SPEAR have greater Moran’s I values than the median of the background. In contrast, SVGs identified by STAMarker and SPARK-X did not show statistically significant differences from the background genes. Here, background genes were defined as all genes excluding common SVGs identified by the four methods. We focused on genes expressed in 150-3,000 spots to optimally capture structured spatial patterns while avoiding sparse or ubiquitous expression profiles (Fig. S4). This comparative advantage against all other methods underscores SEPAR’s enhanced sensitivity in detecting biologically relevant spatial patterns. Thereafter, we compared the SVGs with the 2903 differentially expressed genes identified using DEseq2 [41] by utilizing the manually annotated labels (Fig. 2f). Among the 1,831 SVGs identified by SEPAR, 1,674 were DEGs, significantly surpassing the DEG counts identified by STAMarker (653), SPARK-X (1,337), and SpatialDE (964), demonstrating SEPAR’s superior sensitivity and accuracy in identifying biologically meaningful SVGs. Further, we also compared the SVGs with the highly variable DEGs (defined as the intersection of the DEGs set and the top 3000 HVGs identified using Scanpy package [42] from the original data) (Fig. S5). The SVGs detected by SEPAR show the largest intersection (891 genes), corroborating that SEPAR’s unsupervised SVG detection retains a considerable amount of spatial information. We employed the 1,831 SVGs as input for the spatial clustering methods STAGATE [13] and BayesSpace [12] and compared with their default setting (STAGATE: 3000HVGs, BayesSpace 2000HVGs). As depicted in Fig. S6, utilizing the selected SVGs enhanced clustering performance. The Adjusted Rand Index (ARI) for STAGATE rose from 0.52 to 0.54, and BayesSpace’s ARI improved from 0.46 to 0.52, further affirming the efficacy of the SVGs identified by our method.

SEPAR also demonstrates robust performance in two key downstream applications: gene expression refinement and spatial domain identification. For gene refinement, we selected six marker genes: *CUX2* (a marker for upper layer neurons), *GFAP* (expressed in astrocytes), *SLC17A7* (expressed in excitatory neurons), *RORB* (expressed in layer 4), *PLP1* (a key marker for oligodendrocytes), and *FOXP2* (specifically expressed in deep cortical layers neurons) [43, 44]. Fig. 2g illustrates the original expression and that refined with SEPAR for these six marker genes, where SEPAR effectively reduces noise, thereby enhancing the accurate representation of underlying biological signals. Fig. 2h reveals the differences in Moran’s I distribution for all genes before and after refinement, demonstrating a significant overall increase in Moran’s I. For spatial clustering, we summarize the results of SEPAR, STAGATE [13], GraphST [45], and BASS [14] on 12 DLPFC samples (Fig. S7 and Fig. S8). SEPAR effectively identified the layer structure in the samples, and performs comparably to algorithms specially designed for clustering.

Collectively, applying SEPAR to the DLPFC dataset confirms SEPAR’s robust performance in spatial metagene expression pattern identification, pattern-specific gene analysis, SVG detection, gene expression denoising and spatial domain identification.

### SEPAR identifies distinct laminar structures and spatial patterns in mouse olfactory bulb

We applied SEPAR to analyze the spatial organization of the mouse olfactory bulb using Stereo-seq SRT data [33]. The olfactory bulb exhibits a well-defined laminar architecture, comprising multiple distinct layers from the center outward: the subependymal zone (SEZ), granule cell layer (GCL), internal plexiform layer (IPL), mitral cell layer (MCL), outer plexiform layer (OPL), glomerular layer (GL), olfactory nerve layer (ONL), and meninges. These anatomical layers were previously identified in the original study through spatially constrained clustering (SCC), which integrated both spatial proximity and transcriptomic similarity in a graph-based approach (Fig. 3a) [33]. Our analysis using SEPAR successfully recapitulated these known anatomical structures while providing additional insights into their molecular characteristics.

**Fig. 3:**
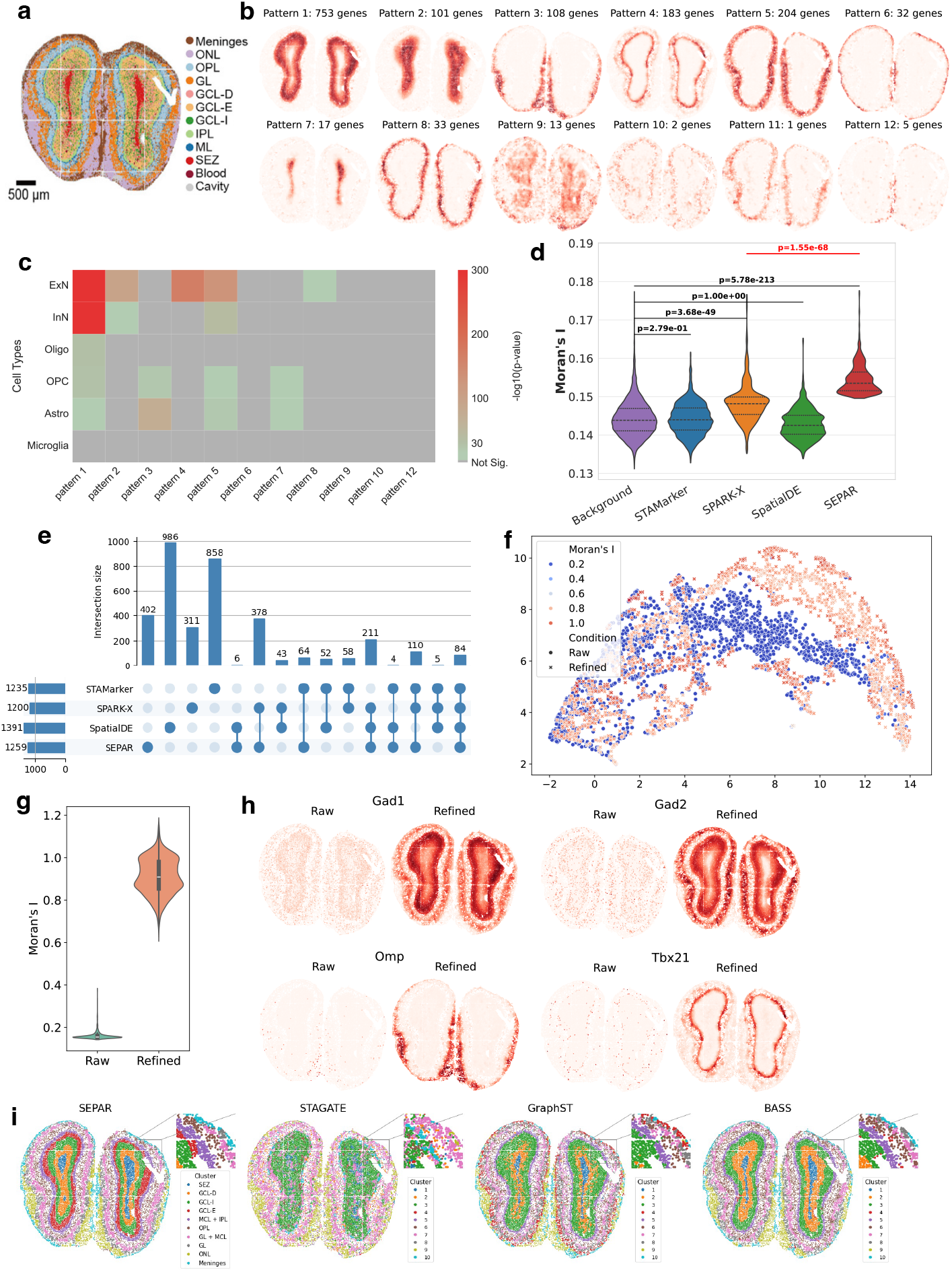
Analysis of the mouse olfactory bulb SRT data generated by Stereo-seq. (**a**) Unsupervised spatially clustering results from original research [33]. (**b**) 12 spatial patterns of metagenes detected by SEPAR with number of pattern-specific genes. (**c**) Cell-type enrichment analysis on the gene sets corresponding to each pattern using CellGO. (**d**) Comparison of Moran’s I values for uniquely identified SVGs by each method. Background genes are all genes excluding the common SVGs identified by the four methods. One-sided Mann-Whitney U test is used to calculate the p-values. (**e**) UpSet plot for SVGs identified by STAMarker, SPARK-X, SpatialDE and SEPAR. (**f** ) UMAP plot of gene expression before and after refinement, colored by Moran’s I and shaped by condition. (**g**) Violin plot for Moran’s I of genes before and after gene expression refinement. (**h**) Visualization of 4 marker genes before and after gene expression refinement. (**i**) Spatial domain identification results of SEPAR, STAGATE, GraphST and BASS.

SEPAR identified 30 spatial metagene expression patterns with the 12 most significant ones shown in Fig. 3b. The number of pattern-specific genes ranges from 5 to 753, and these patterns revealed distinct laminar structures. We then performed cell-type enrichment analysis using CellGO and GO term enrichment using g:Profiler for the pattern-specific gene sets (Fig. 3c and Fig. S9). The gene set for Pattern 7 is closely linked to processes such as “regulation of neuron differentiation” (GO:0045664, p-value = 2.97e-04) and “neurogenesis” (GO:0022008, p-value = 4.53e-04), both of which are critical to the SEZ, a recognized neurogenic niche. The enrichment of “oligodendrocyte differentiation” (GO:0048709, p-value = 1.21e-03) supports the cell-type enrichment results showing astrocytes and oligodendrocyte precursor cells are abundant in this region, consistent with its role in ongoing neural development and repair. Pattern 6, meanwhile, is characterized by terms like “collagen-containing extracellular matrix” (GO:0062023, p-value = 3.76e-11) and “extracellular matrix structural constituent conferring compression resistance” (GO:0030021, p-value = 8.75e-05). The spatial location of Pattern 6 aligns with the meninges as identified in prior research [33], demonstrating that SEPAR is a reliable tool for unsupervised identification of biologically significant regions based on spatial distribution of decomposed patterns and subsequent gene-specific analysis.

According to the metagene patterns, SEPAR identified 1,259 SVGs (Fig. 3e). Compared with STAMarker, SPARK-X, and SpatialDE, 84 genes were identified in common, with SEPAR and SPARK-X sharing the largest overlap (378 genes), and SEPAR and STAMarker sharing the second largest overlap (64 genes). To evaluate the spatial significance of SVGs uniquely identified by each individual method, the Moran’s I values were compared (Fig. 3d). Compared to the background genes, the SVGs uniquely identified by SEPAR and SPARK-X exhibited significantly higher Moran’s I values (p-value = 5.78e-213 and p-value = 3.68e-29, respectively, one-sided Mann-Whitney U test), with SVGs identified by SEPAR showing significantly higher values than those using SPARK-X (p-value = 1.55e-68). The SVGs identified by SEPAR have higher Moran’s I values than the median of the background genes. In contrast, the SVGs identified using STAMarker and SpatialDE did not show statistically significant difference from the background genes, which were defined as all genes excluding the common SVGs identified by the four methods. Here genes expressed in 100-5000 spots were considered (Fig. S10). Consistent with our previous findings, SEPAR demonstrated superior statistical significance across all compared methods, further confirming its enhanced capability to detect spatially structured gene expression patterns across different biological contexts.

Gene expression denoising using SEPAR demonstrated significant improvement in spatial smoothness. Fig. 3f presents the UMAP visualization of gene expression before and after refinement, showing minimal changes in distribution, indicating that the adjustments did not distort the original data significantly. Additionally, as shown in Fig. 3g, the spatial autocorrelation of gene expression increased substantially after refinement. We further examined several marker genes: *Gad1* and *Gad2* (Glutamate Decarboxylase 1 and 2) for GABAergic neurons, *Omp* (Olfactory Marker Protein) for mature olfactory sensory neurons, and *Tbx21* (*T-box 21*) for mitral and tufted cells [43, 46]. The comparison of pre- and post-denoising in Fig. 3h demonstrates a marked enhancement in the spatial structure, particularly for *Tbx21*, whose post-denoising pattern aligns well with the mitral cell layer (MCL) of the olfactory bulb, predominantly composed of Mitral Cells.

Finally, we compared SEPAR’s clustering performance with STAGATE, GraphST, and BASS (Fig. 3i). SEPAR’s clustering results closely resemble those of GraphST and BASS, while demonstrating superior definition of laminar structures. In contrast, STAGATE’s clustering results were less distinct for this dataset.

### SEPAR enables efficient spatial clustering and expression refinement in image-based spatial transcriptomics

Image-based spatial transcriptomics technologies that utilize fluorescence in situ hybridization or in situ sequencing, such as seqFISH+ [47], osmFISH [34], MER-FISH [4], and STARmap [48], can offer single-cell resolution expression and precise spatial information of RNA transcripts. These methods utilize pre-designed probes and high-resolution microscopy for detection, thus enabling absolute quantification of transcripts with spatial context, but they are restricted to limited pre-selected genes compared to sequencing-based spatial transcriptomics methods. Given these characteristics, we emphasize spatial domain identification and gene expression refinement when using SEPAR for downstream analysis. We applied SEPAR to mouse somatosensory cortex dataset generated by osmFISH [34] and mouse hypothalamic preoptic region dataset generated by MERFISH [4].

We first analyzed the osmFISH dataset from the mouse somatosensory cortex [34], which comprises only 33 genes. Fig. 4a shows the spatial domains identified by different methods. SEPAR achieved superior results, with an ARI of 0.666 and an NMI score of 0.703, significantly outperforming STAGATE (ARI = 0.387), GraphST (ARI = 0.314) and BASS (ARI = 0.429). Since SEPAR is developed based on matrix factorization, the model training process is much faster than the neural network-based methods. As illustrated in Fig. 4b, SEPAR exhibits the shortest runtime among the four methods evaluated, requiring only 6.3 seconds – about one-quarter of the runtime of the second-fastest method GraphST.

**Fig. 4:**
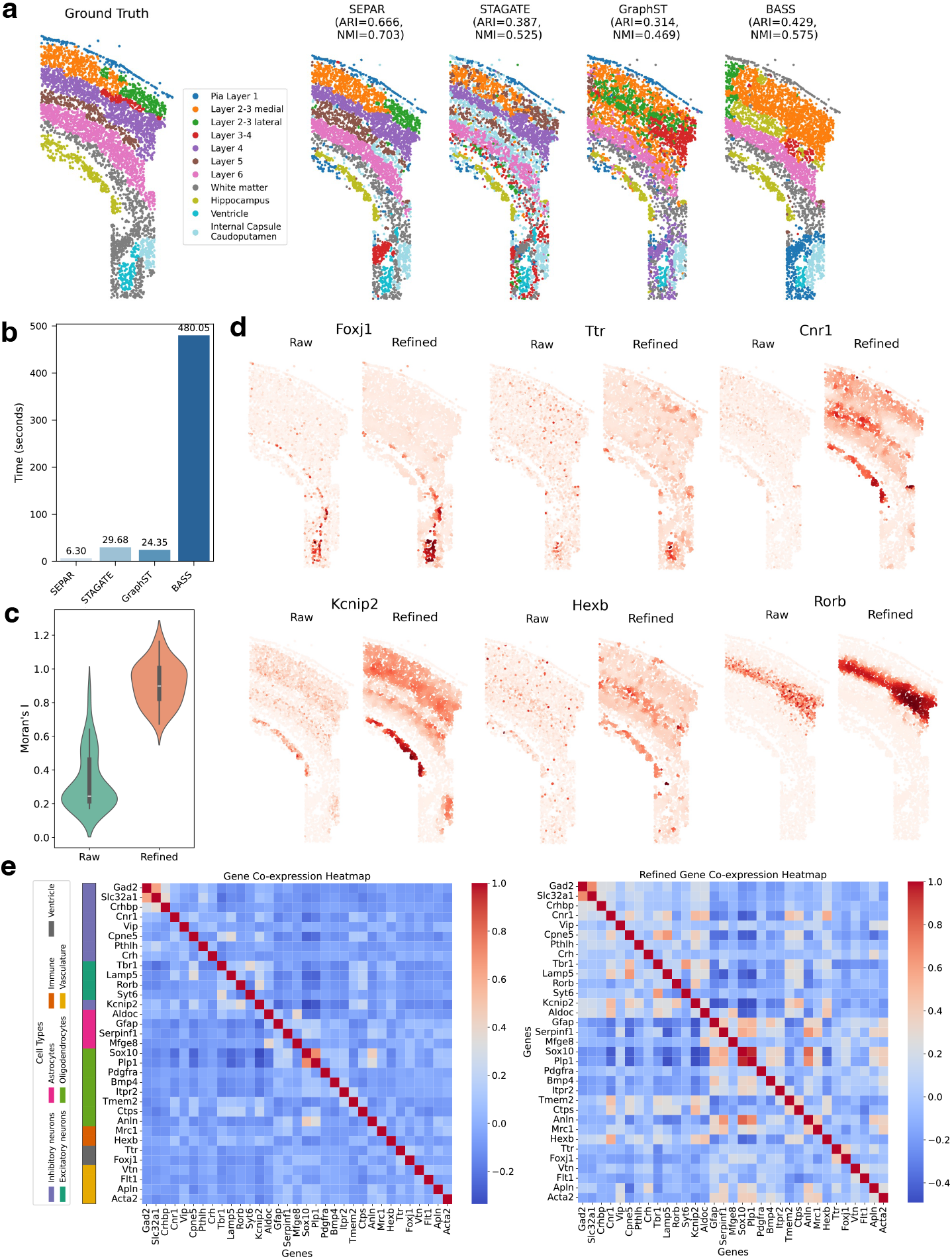
Analysis of the mouse somatosensory cortex data generated by osmFISH. (a) Spatial domain identification using SEPAR, STAGATE, GraphST and BASS. (b) Comparison of running time for SEPAR, STAGATE, GraphST and BASS. (**c**) Violin plots of Moran’s I values for genes expression before and after refinement. (**d**) Visualization of 6 genes before and after gene expression refinement. (**e**) Gene co-expression heatmaps generated from raw (left) and refined (right) expression data, with cell-type markers annotated based on original osmFISH study [34].

Next, we performed gene refinement and spatial gene co-expression analysis, demonstrating that gene refinement helps detect more correlated genes. As observed in Fig. 4d, gene refinement notably reduced noise, enhancing cortical structure clarity and significantly improving Moran’s I values of the genes (Fig. 4c). After gene refinement, more spatially correlated genes are identified (Fig. 4e): 23 pairs showed strong correlations (correlation coefficient *>* 0.4) after refinement, whereas only 3 pairs were observed in the raw data. As an example, genes *Foxj1* and *Ttr* show much higher correlations after gene refinement (Fig. 4d). According to the existing research [34], both *Foxj1* and *Ttr* are marker genes for ependymal and choroid plexus cells, respectively. These cells are prominently distributed in the ventricle, showing anatomical adjacency and functional interactions. As another example, *Cnr1, Kcnip2* and *Hexb* show much clearer correlated patterns after applying SEPAR (Fig. 4d,e). Both *Cnr1* and *Kcnip2* relate to inhibitory neurons, while *Hexb* is associated with microglial immune cells. While the spatial pattern of *Hexb* is not visually apparent in Fig. 4d, violin plots of raw *Hexb* expression across spatial domains (Fig. S12) confirm its elevated expression in the hippocampus and Layer 5, suggesting its role in neural plasticity and immune function. Despite having only 33 genes in the osmFISH dataset, SEPAR demonstrated its capability for effective analysis.

To further validate the performance of SEPAR, we analyzed a MERFISH [4] dataset from the mouse hypothalamic preoptic region [35], which measured the expression of 155 genes across 5926 cells. We directly used the spatial patterns extracted using SEPAR as low-dimensional representations for clustering. As shown in Fig. 5a and Fig. 5b, the computation of SEPAR is much faster than BASS, though the spatial domain identification accuracy of SEPAR ranks the second among four methods, just behind BASS. Given that the ground truth annotations were originally provided by BASS [14], SEPAR’s competitive performance (ARI = 0.454, NMI = 0.565) demonstrates its robust capability in spatial domain identification. Notably, when comparing with the Allen Brain Atlas [49], SEPAR’s identification of the medial preoptic nucleus (MPN) actually showed better anatomical correspondence than BASS (Fig. S10), suggesting its biological relevance. SEPAR costs only 9.13 seconds for the computation, while BASS takes 508.77 seconds. In contrast, STAGATE and GraphST reached lower spatial domain identification accuracy with more computational time (STAGATE: 43.03s, GraphST: 21.73s). SEPAR demonstrates the highest computational efficiency, balancing efficacy and efficiency among the four methods.

**Fig. 5:**
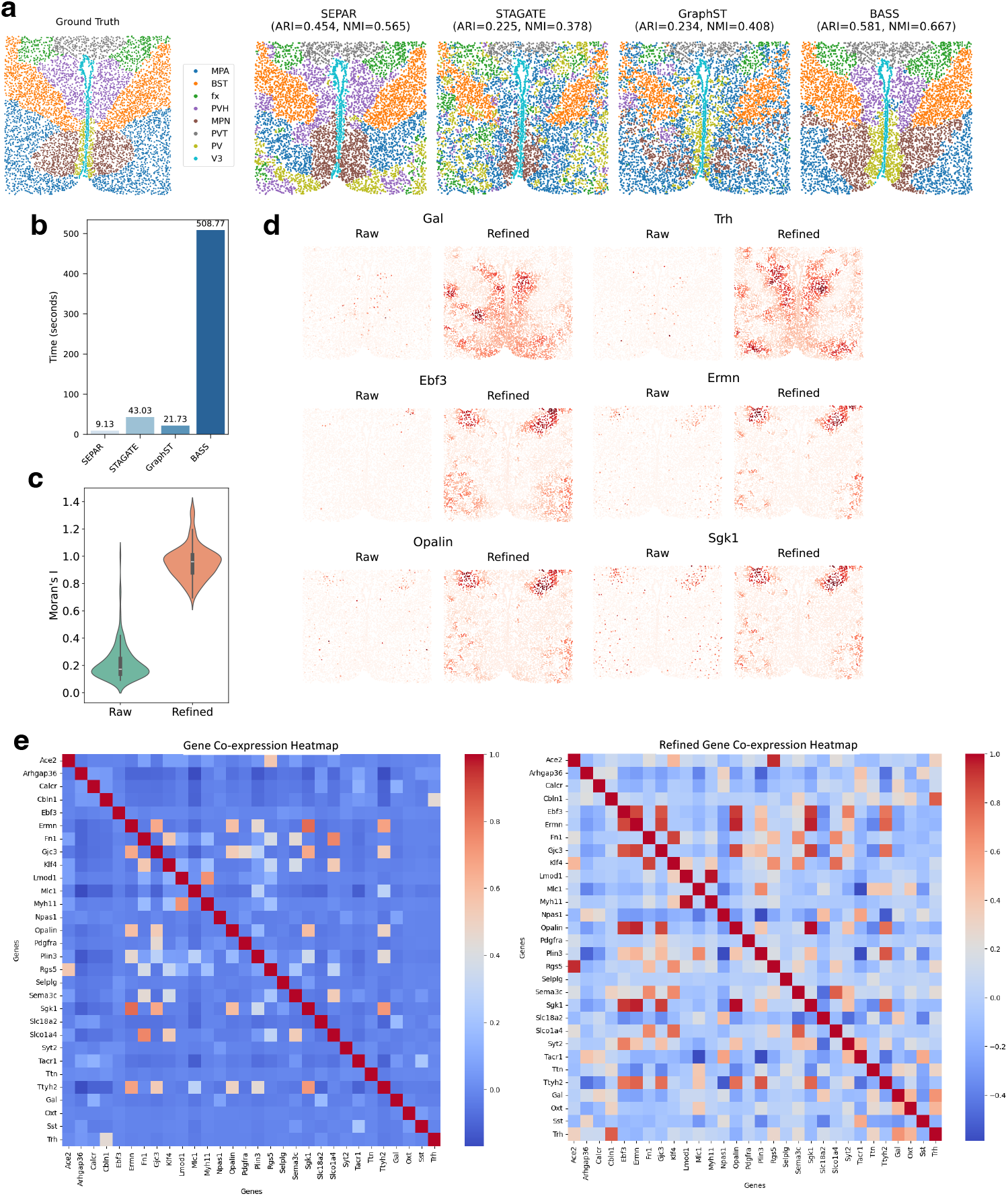
Analysis of the mouse hypothalamic preoptic region generated by MERFISH. (a) Spatial domain identification using SEPAR, STAGATE, GraphST and BASS. The ground truth annotation was obtained from [14]. (**b**) Comparison of running time for SEPAR, STAGATE, GraphST and BASS. (**c**) Violin plots of Moran’s I values for gene expression before and after refinement. (**d**) Visualization of 6 genes before and after gene expression refinement. (**e**) Gene co-expression heatmaps generated from raw (left) and refined (right) gene expression.

We also conducted gene expression refinement and spatial gene co-expression analysis on this dataset. After the refinement, the spatial correlation of gene expression is significantly improved (Fig. 5c). Fig. 5d displays the expression levels of 6 genes before and after refinement, showing enhanced spatial patterns. Fig. 5e presents a co-expression heatmap of the top 30 highly variable genes before and after refinement, 31 pairs showing strong correlations (correlation coefficient *>* 0.5) after refinement, whereas only 15 correlations were observed in the raw data. The refined data not only exhibited correlation improvement of known correlated genes but also identified new correlated gene pairs. For instance, *Gal* and *Trh* were not recognized as co-expressed genes in the raw data, while they showed highly correlated expression patterns in similar domains after refinement, with corresponding cells being spatially proximate, as previously shown in Fig. 5d. Similarly, *Ebf3, Ermn, Opalin*, and *Sgk1*, whose spatial expression patterns are visualized in Fig. 5d, were identified as correlated gene pairs after refinement, with their high-expression cells exhibiting clear spatial adjacency in the tissue. This illustrates that gene refinement using SEPAR enables spatial gene co-expression analysis, leveraging spatial location information.

### SEPAR identifies the associations between genes and proteins in spatial CITE-seq data

Recent technological advances [10, 11] have enabled simultaneous profiling of multiple molecular modalities such as RNA, protein, and chromatin accessibility within the same spatial context, offering unprecedented opportunities to study the complex interplay between different molecular layers. We next explored the capability of SEPAR in vertical integration of multi-omics data with two datasets generated by spatial CITE-seq and MISAR-seq, respectively.

First, SEPAR was applied to a spatial CITE-seq dataset from human tonsil tissue [10]. Spatial CITE-seq enables simultaneous measurement of gene expression and protein levels, providing complementary molecular information. Here, the metagenes are extended to meta-gene-proteins. Six spatial patterns corresponding to meta-gene-proteins identified are displayed in Fig. 6a. Among these, four patterns: Pattern 1, 2, 4, and 5 incorporate cross-omics information. We hypothesized that these spatially matched gene-protein sets harbor potential gene-protein relationships. To validate this hypothesis, we selected the four most significant genes and proteins from Pattern 1, 2, 4, and 5, and generated a co-expression heatmap with denoised expression level (Fig. 6b). The analysis revealed higher correlation coefficients between genes and proteins within the same pattern (mean = 0.622) compared to those between different patterns (mean = − 0.086). Despite the inherent heterogeneity between protein and gene data, SEPAR successfully extracted shared patterns, linking RNA with protein data. For example, the co-expression between CD74 and *HLA-C* identified in our analysis is consistent with established biological mechanisms, where CD74 has been shown to regulate both MHC class II and certain MHC class I molecules in antigen 15 presentation [50]. Furthermore, enrichment analysis was performed on the identified pattern-specific genes and proteins, with results for Pattern 4 shown in Fig. 6c. The enrichment analysis revealed significant immune-related functions, particularly in cell surface components and immune response pathways. Notably, the enrichment of “external side of plasma membrane” and “cell surface” components, along with immune response pathways and hematopoietic cell lineage, indicates robust immune cell activities and extensive cell-cell interactions in the tissue microenvironment.

**Fig. 6:**
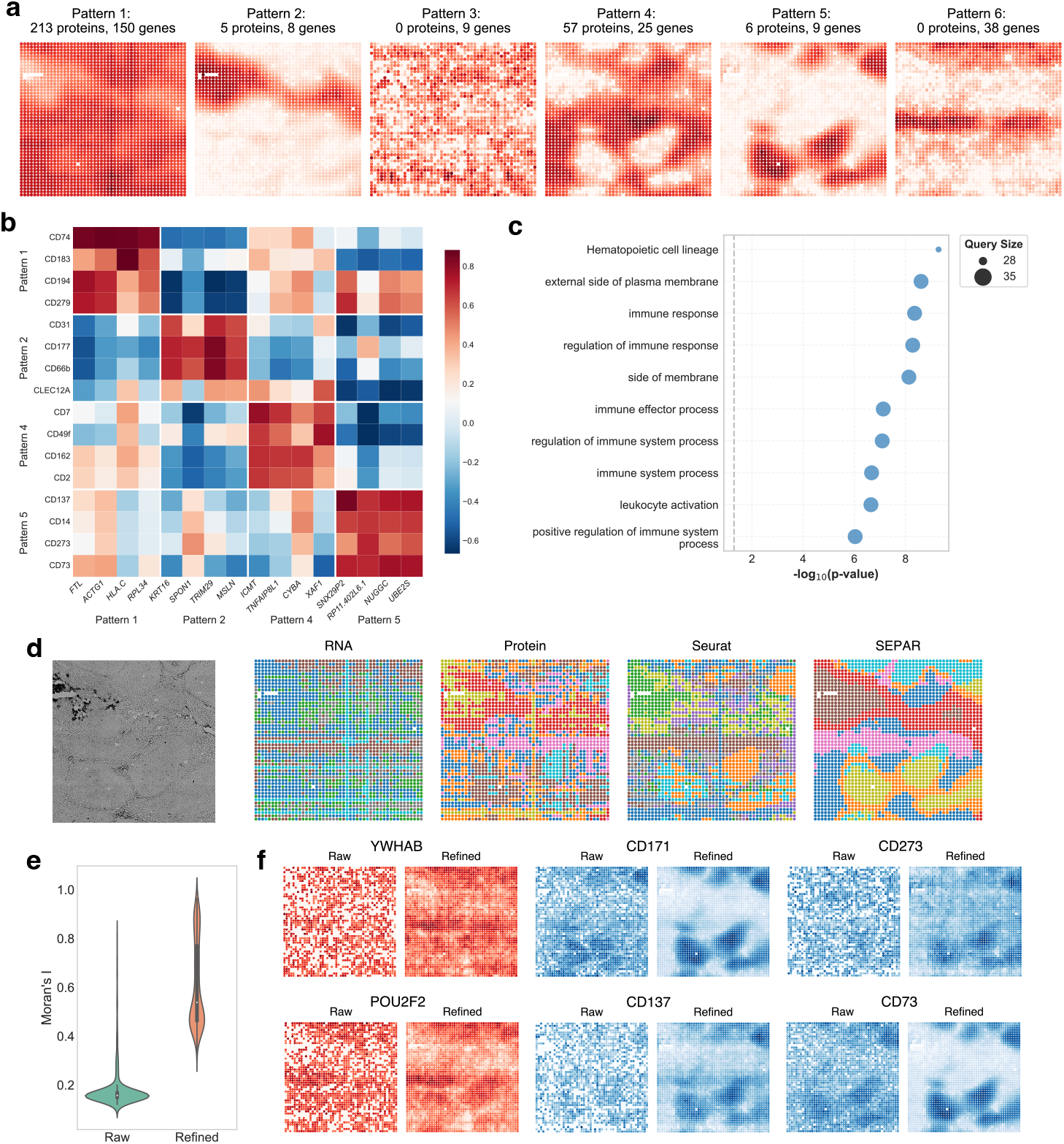
Integrative analysis of spatial CITE-seq data from human tonsil tissue using SEPAR. (**a**) Six spatial patterns identified by SEPAR within the spatial CITE-seq dataset, detailing the associated proteins and gene counts for each pattern. (**b**) Co-expression heatmap with denoised expression of selected genes and proteins from Pattern 1, 2, 4, and 5. (**c**) Enrichment analysis for Pattern 4 showing significant biological processes and pathways. (**d**) Comparison of spatial clustering between SEPAR and Seurat. High-resolution microscope images (leftmost) provide reference for tissue architecture, followed by Seurat’s clustering results from individual modalities (RNA-based, protein-based, resolution 0.8), Seurat’s integrated RNA+protein analysis (WNN-based, resolution 1.2), and SEPAR. (**e**) Violin plots of Moran’s I value for genes before and after refinement. (**f** ) Visualization of 2 genes (red) and 4 proteins (blue) before and after refinement.

The spatial patterns corresponding to meta-gene-proteins identified using SEPAR also enabled unsupervised spatial clustering and expression refinement, with comparative clustering results shown in Fig. 6d. The figure presents four clustering approaches: SEPAR’s integrated analysis and Seurat’s individual analyses of RNA, protein, and combined RNA+protein data using weighted nearest neighbor (WNN) framework. SEPAR’s clustering results exhibited enhanced spatial coherence with well-defined domain boundaries, effectively capturing the underlying tissue architecture. In contrast, the individual modality analyses and even the combined RNA+protein approach from Seurat showed more fragmented clustering patterns. This comparison demonstrates SEPAR’s capability in leveraging cross-omics information to achieve more biologically meaningful spatial domain identification.

As for the refinement effect, quantitative assessment using Moran’s I statistics demonstrated significant enhancement in spatial coherence for both gene and protein expressions (Fig. 6e). The refinement process substantially improved data quality (Fig. 6f), where the initially noisy and ambiguous gene expression patterns were transformed into well-defined spatial distributions with intensified characteristic features. This enhanced data quality enabled comprehensive spatial co-expression analysis of the top 15 highly variable genes and proteins (Fig. S13), revealing markedly improved molecular correlations: 44 pairs of biological molecules showed strong correlations (correlation coefficient *>* 0.5) after refinement, whereas no such strong correlations were observed in the raw data. For example, we observed co-expression patterns between genes *YWHAB* and *POU2F2*, and the proteins CD71, CD137, CD73, and CD273 (Fig. 6f), demonstrating SEPAR’s ability to detect cross-modality relationships after refinement.

### SEPAR identifies the associations between gene expression and ATAC in MISAR-seq data

We also applied SEPAR directly for integrating spatial ATAC-seq and SRT data on a mouse embryonic (E15.5) brain dataset obtained through MISAR-seq [11]. MISAR-seq is a novel technology that simultaneously profiles chromatin accessibility and gene expression within the spatial context, providing insights into gene regulation mechanisms. The 14 significant spatial patterns corresponding to meta-gene-peak identified are displayed in Fig. 7a, with 11 patterns linked to both ATAC and RNA data, indicating SEPAR’s capability in effectively connecting RNA with chromatin accessibility. To validate these connections, the genes associated with the pattern-specific peaks were first identified using ChIPseeker [51] with those absent from the gene expression data deleted. This gene set was then compared with the previously identified pattern-specific genes. The comparison revealed substantial overlaps: in Pattern 1, 12 out of 17 genes identified from the pattern-specific peaks were found in the pattern-specific gene set, while in Pattern 4, 8 genes overlapped between 26 pattern-specific genes and 20 genes obtained from pattern-specific peaks (Fig. 7b), suggesting strong biological relevance between identified pattern-specific genes and peaks. We selected two most significant pattern-specific genes and peaks from each of the 10 patterns and generated a co-expression heatmap with denoised expression level (Fig. 7c). The analysis revealed higher correlations between genes and peaks within the same pattern (mean = 0.613) compared to those between different patterns (mean = 0.028), demonstrating the expression-level concordance of genes and peaks identified using SEPAR.

**Fig. 7:**
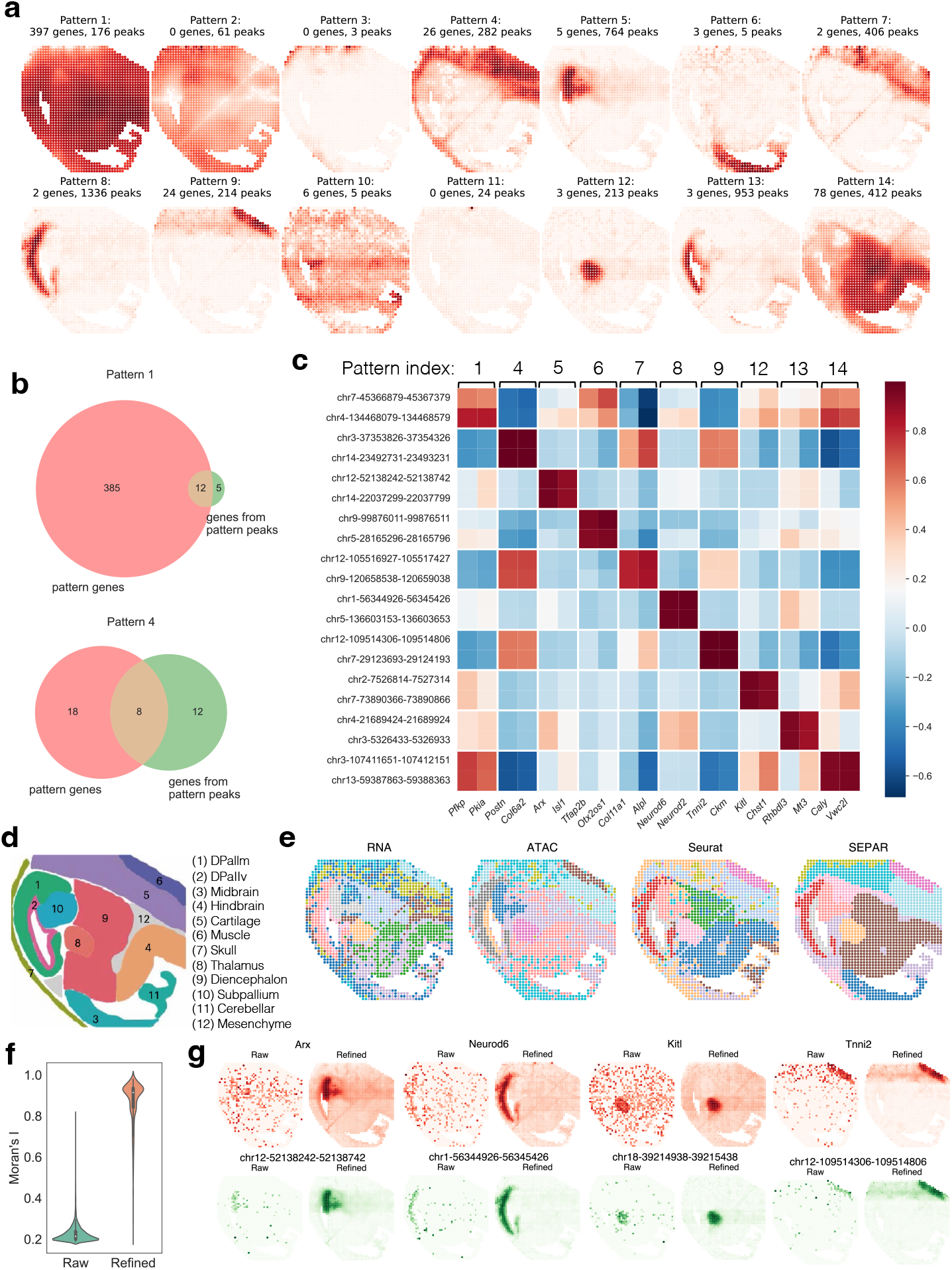
Analysis of MISAR-seq data from mouse embryonic (E15.5) brain tissue using SEPAR. (**a**) Fourteen spatial patterns identified by SEPAR within the MISAR-seq dataset, detailing the associated peaks and gene counts for each pattern. (**b**) Gene-peak association analysis showing overlap between pattern-specific gene sets and pattern-specific peaks’ associated genes. (**c**) Co-expression heatmap with denoised expression of selected genes and peaks from identified spatial patterns. (**d**) Reference annotation map of mouse embryonic brain regions. (**e**) Comparison of spatial clustering between SEPAR and Seurat. Left to right: clustering results from individual modalities (RNA-based, ATAC-based) using Seurat, Seurat’s integrated RNA+ATAC analysis, and SEPAR clustering. (**f** ) Violin plots of Moran’s I values for gene expression before and after refinement. (**g**) Visualization of four genes (red) and four peaks (green) before and after refinement.

As for the spatial domains, compared with the 11 annotated spatial domains in the mouse embryonic (E15.5) brain (Fig. 7d), SEPAR accurately identified key brain regions including midbrain and thalamus, as well as surrounding non-neural tissues like muscle (Fig. 7e). Notably, SEPAR achieved more coherent spatial domains with reduced noise and enhanced structural definition compared to Seurat’s individual analyses of RNA, ATAC, and combination of RNA and ATAC.

The refinement process led to significantly improved spatial autocorrelation as demonstrated by Moran’s I statistic (Fig. 7f). Spatial co-expression analysis of the top 15 highly variable genes and peaks (Fig. S14) revealed substantially improved molecular relationships: 137 pairs showed strong correlations (correlation coefficient *>* 0.5) after refinement, whereas no such correlations were observed in the raw data. Several notable co-expression pairs were identified, for example, genes *Col1a2, lgf2, Airn* with their corresponding peaks chr10-80260918-80261418, ch8-123411167-123411667, and chr7-141949425-141949925 (Fig. 7g). These multi-omics pairs, which share similar spatial patterns and were previously masked by noise in the raw data, became apparent after refinement. Additionally, analysis of spatially variable genes and peaks (Fig. S15, S16) corroborated the identified cross-omics spatial patterns.

To further characterize the regulatory mechanisms underlying these spatial patterns, we performed motif enrichment analysis for all pattern-specific peaks using HOMER (Hypergeometric Optimization of Motif EnRichment) [52] (Fig. S17). This analysis revealed distinct transcription factor binding motifs enriched in different spatial domains. Pattern 8, localized in the developing pallium (DPallm), showed enrichment of proneural basic helix-loop-helix (bHLH) transcription factor binding sites, including NeuroD1, NeuroG2, and Atoh1. For this pattern, we identified only two pattern-specific genes: *Neurod2* and *Neurod6*, both of which are essential for proper neuronal migration and differentiation [53]. The spatial co-occurrence of NeuroD1 and NeuroG2 binding motifs and the expression of Neurod2 and Neurod6 in the developing pallium aligns with their known roles in neuronal differentiation [53, 54], though their regulatory relationships remain to be determined. Pattern 9, localized in the developing muscle tissue surrounding the E15.5 mouse brain, exhibited highly significant enrichment of E-box motifs (CANNTG) (Fig. S14), which are canonical binding sites for myogenic regulatory factors (MRFs) [55]. The most significantly enriched motifs were from the bHLH family of myogenic regulatory factors, with Myf5 showing the strongest enrichment (p = 1e-66, 78.30% of target sequences), followed by MyoD (p = 1e-59, 78.77%) and MyoG (p = 1e-56, 85.85%). This pattern’s specific genes comprised a comprehensive set of muscle structural components, including myosin heavy chains (*Myh3, Myh8*), troponins (*Tnnt1, Tnnt3, Tnni1, Tnni2, Tnnc2*), and other essential sarcomeric proteins (*Acta1, Actc1, Des, Ttn, Mybpc1, Neb*). The extensive presence of MRF binding motifs in pattern-specific peaks, together with these muscle-specific genes, suggests active regulatory relationships in this muscle domain. Notably, TCF21 binding motifs showed strong enrichment (p = 1e-59, 83.02% of target sequences) in pattern-specific peaks, along with TCF12 (p = 1e-52, 82.08%). TCF12 (also known as HEB) preferentially expresses in developing muscle and forms heterodimers with MyoD to regulate muscle-specific genes [56, 57], while TCF21 directly binds to regulatory regions of both Myf5 and MyoD to function as their upstream activator [58]. The presence of these functionally diverse but muscle-related genes, along with their experimentally validated regulatory interactions, demonstrates the potential of SEPAR’s multi-omics pattern-specific analysis in identifying comprehensive tissue-specific regulatory networks.

## 3 Discussion

In this work, we introduced SEPAR, a robust method designed to identify the spatial patterns of metagene expression along with the associated pattern-specific gene sets for SRT data. SEPAR is a graph-regularized NMF-based approach that leverages the non-negative nature of gene expression data and NMF’s interpretability to uncover the underlying patterns and structures for subsets of genes in SRT data. Using the identified spatial metagene expression patterns, SEPAR excels in various downstream analytical tasks, including spatial pattern-specific gene analysis, SVG identification, spatial clustering, and gene expression denoising. Unlike traditional methods for spatial domain identification that focus on the expression pattern of all genes, SEPAR naturally groups the genes into different patterns corresponding to the metagenes. This holistic approach allows SEPAR to identify genes with better cell-type enrichment and gene ontology enrichment and significant regional differential expression without any supervision.

A key advantage of SEPAR is its versatility across different data types. Beyond demonstrating robust performance in diverse spatial transcriptomics technologies (10 × Visium, Stereo-seq, osmFISH and MERFISH), where SEPAR consistently identified biologically meaningful spatial patterns and revealed tissue-specific expression programs across different resolution scales and measurement principles, SEPAR effectively handles multi-omics spatial molecular data. It successfully uncovered coordinated spatial patterns between RNA and protein expression in spatial CITE-seq data, as well as those between chromatin accessibility and gene expression in MISAR-seq data, revealing strong correlations between co-localized molecules and mechanistic insights into spatial gene regulation.

The success of SEPAR in spatial pattern identification and downstream analysis for one tissue slice has motivated us to extend it to simultaneously analyze multiple slices of tissue SRT data from the same sample source, which is named SEPARmult (**Supplementary Note 1**). Experiments on the 12 tissue slices from the DLPFC data show that in spatial domain identification across all the samples, SEPARmult significantly improved the median ARI of SEPAR from 0.520 to 0.590, outperforming other state-of-the-art methods including STAGATE (median ARI = 0.525), GraphST (median ARI = 0.515), and BASS (median ARI = 0.465). This extension validates the scalability and adaptability of SEPAR to more complex analytical scenarios.

While SEPAR is applicable to various scenarios, the increasing generation of spatial data presents several promising research directions. First, incorporating single-cell RNA sequencing data and cell-type annotations could provide deeper biological insights. This integration could bridge the gap between spatial domains and cellular composition, enabling simultaneous domain identification and cell-type deconvolution while leveraging the rich cell-type information from single-cell references. Second, the flexible framework of SEPAR creates opportunities for incorporating biological prior knowledge. For instance, integrating gene regulatory networks could help identify coordinated spatial patterns of transcription factors and their targets, potentially revealing tissue-specific regulatory mechanisms and developmental programs in their spatial context. Third, the principles underlying SEPARmult’s success in multi-slice integration could be extended to cross-platform integration of spatial transcriptomics data. This remains a critical challenge in the field, as different platforms exhibit distinct technical characteristics and batch effects, yet integrating data across platforms could significantly increase sample sizes and statistical power while enabling cross-validation of findings.

## 4 Methods

### Spatial metagene expression pattern recognition model

We developed a novel graph regularized non-negative matrix factorization (NMF)- based model to conduct Spatial metagene Expression PAttern Recognition (SEPAR). The model is formulated as:

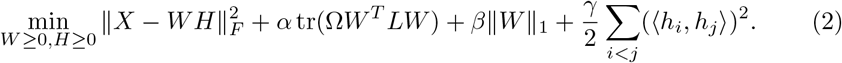

Here, *X* is the *n* × *p* gene expression matrix corresponding to *n* spots (cells) and *p* genes. *W* ={ *w*_1_, *w*_2_, … , *w*_*r*_} represents the *r* spatial metagene expression patterns, and each *h*_*i*_ of *H* = {*h*_1_, *h*_2_, … , *h*_*r*_ }^*T*^ indicates the *i*-th metagene expression across the *p* genes. *L* is the graph Laplacian matrix incorporating the spatial location and gene expression information. The weight matrix Ω is taken to be the diagonal matrix of Pattern Significance Scores (diag(*PSS*)), which measures the influence of the graph regularization for spatial metagene expression patterns. *α, β*, and *γ* are parameters controlling the weight of each regularization term. The weighted graph penalty term ensures that close spots have similar pattern representations. The sparsity penalty in helps identify distinct and localized spatial patterns. And the final term promotes orthogonality between the metagenes *h*_*i*_ and *h*_*j*_ in matrix *H*, and minimizing this term reduces redundancy, guaranteeing pattern distinctness.

#### Graph construction

An undirected graph is built based on a predefined radius *r*^*^or a computed radius:

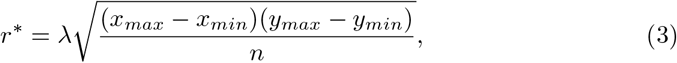

where *x*_*max*_, *x*_*min*_, *y*_*max*_, and *y*_*min*_ represent the maximum and minimum values of the 2D coordinates for all spots/cells, and *λ* is a scaling parameter, typically set to *λ* = 1.1. The adjacency matrix *A* of the graph is defined as:

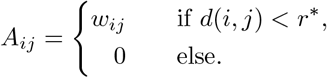

The weight *w*_*ij*_ for the edge between spots *i* and *j* is determined by the cosine distance *d*_*cos*_(*i, j*):

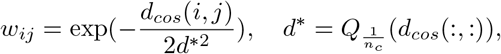

where 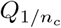 is the 1*/n*_*c*_ quantile, and *n*_*c*_ is the estimated number of clusters in the data. The Laplacian matrix *L* is then computed as *L* = *D*− *A*, where *D* is a diagonal matrix with the diagonal entries being the degree of the corresponding spot.

#### Pattern significance score (PSS)

To quantitatively evaluate the biological relevance and spatial coherence of extracted metagene expression patterns, we developed a metric PSS to measure their significance. For any given pattern *i*, we identified its corresponding spatial distribution *w*_*i*_ and the coefficient *h*_*i*_ for each gene. The pattern significance score (PSS) is defined as:

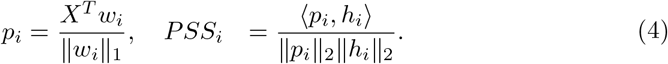

A higher value of *PSS*_*i*_ indicates that the *i*-th gene expression pattern represented by *w*_*i*_ predominantly occurs in the spatial locations.

We iteratively solve the optimization problem (2). Following the multiplicative update framework, we derive the following update rules that guarantee non-negative solutions and monotonically decreasing of the objective function:

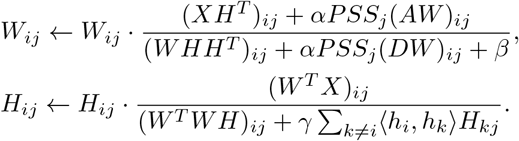

The whole computation procedure is summarized in Algorithm 1.

#### Multi-omics analysis

For integrated analysis of multiple data modalities (RNA, ATAC-seq, protein) from the same spatial locations, after we implement preprocessing for all the modalities, the processed data matrices are concatenated to form an integrated matrix *X* = [*X*_*RNA*_, *X*_*AT* *AC*_, *X*_*protein*_]. Then, SEPAR is applied to this integrated matrix to identify shared spatial patterns across modalities.

##### Algorithm 1 Spatial gene expression pattern recognition algorithm

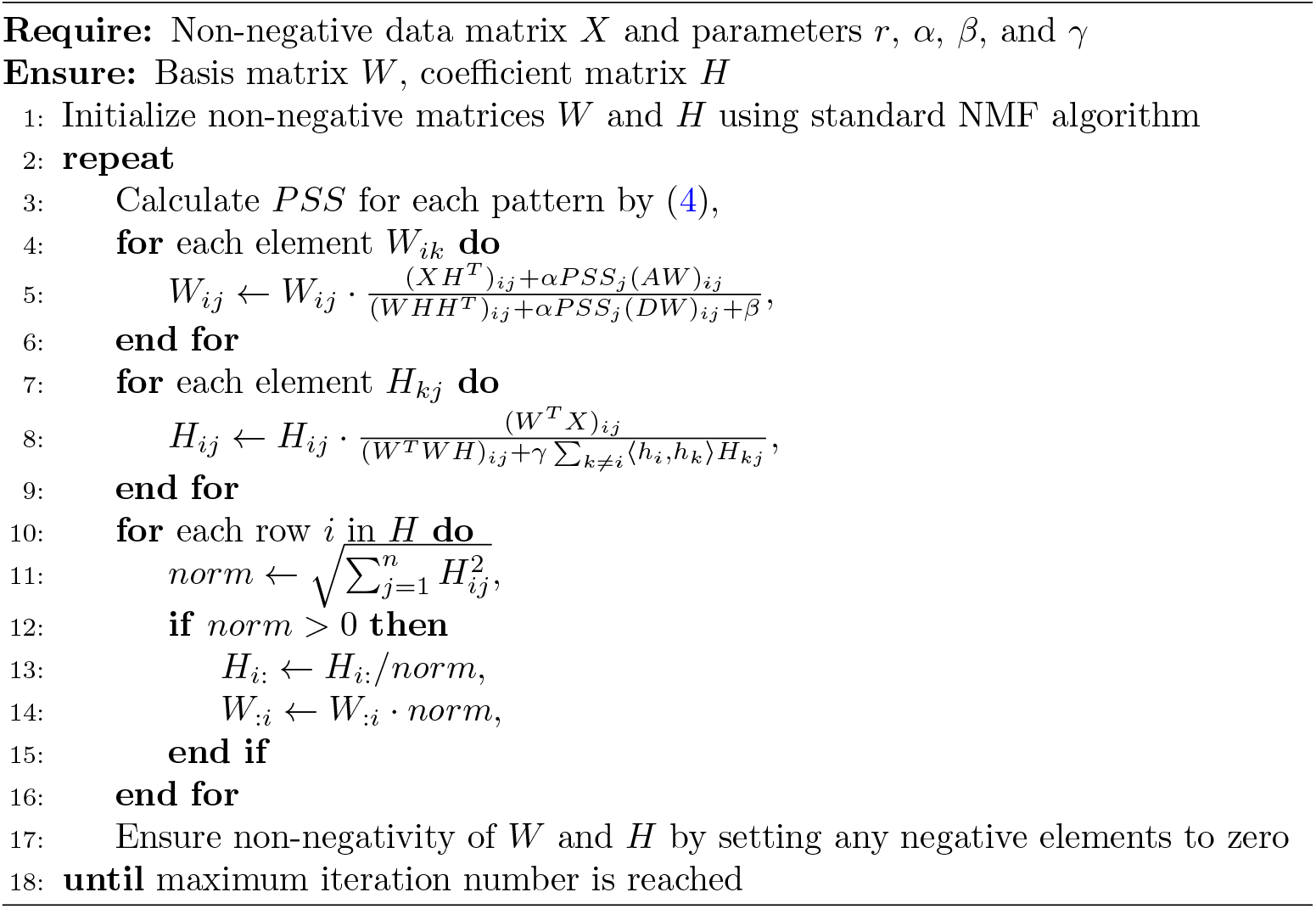

### Downstream tasks

#### Pattern-specific gene detection

To identify the genes specifically associated with each spatial metagene expression pattern, we implement a two-step normalization process on the matrix *H*:

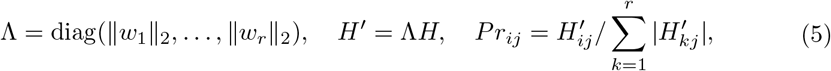

where *Pr* represents the normalized pattern contribution matrix to the genes. Each column of *Pr* quantifies the relative contribution of all patterns to a specific gene. A gene is classified as pattern-specific when its corresponding entry in *Pr* exceeds a threshold *ϵ*_*_, indicating strong association with a particular spatial metagene expression pattern. **Selection of SVGs** SVGs are different from the pattern-specific genes in that they can be expressed on more than one metagene expression pattern. The SVGs are identified by evaluating the contribution of each spatial pattern to the gene expression. For each gene *i*, we compute the reconstruction error from the combination of identified patterns:

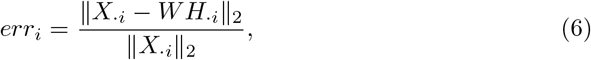

Genes with *err*_*i*_ *< ϵ* are classified as SVGs, indicating their expression exhibits strong spatial organization that aligns with the discovered tissue architecture.

#### Spatial domain identification

The decomposed basis matrix *W*∈ ℝ_*n×r*_ from SEPAR can be taken as a lower-dimensional representation for the spots or cells. Spatial domain identification is performed with *W* . While SEPAR is robust to the choice of dimensionality parameter *r* (typically set to 30 in practice), not all identified patterns necessarily represent biologically meaningful spatial organizations. Indeed, SEPAR’s ability to detect novel spatial patterns beyond conventional anatomical structures (e.g., non-canonical patterns that deviate from classical laminar organization) necessitates a systematic filtering approach to distinguish genuine biological signals from technical artifacts. Therefore, spatial domain identification is performed with *W* through a sequential pattern filtering process. Initially, we filter out *N*_1_ patterns with the smallest *ℓ*_2_-norms, which typically represent low abundance patterns. We further exclude *N*_2_ patterns with the lowest PSS’s to eliminate the potentially noisy components. The resulting filtered low-dimensional matrix *W*^*^ is then subjected to *k*-means clustering to identify distinct spatial domains.

#### Gene expression denoising

The denoised expression data is directly derived from the reconstructed gene expression matrix *WH*. This reconstruction captures the essential low-dimensional structure of the data, effectively filtering out noise.

### Data preprocessing and parameter settings

For data preprocessing, we employed distinct protocols for different spatial transcriptomics platforms. For sequencing-based and multi-omics datasets, we performed log-transformation and library size normalization using the package ‘Scanpy’ [42]. For multi-omics datasets, each modality was processed independently following this protocol. We selected the top 3,000 genes with highest spatial autocorrelation (measured by Moran’s I statistic) for downstream analysis. The sequencing-based spatial transcriptomics data were processed by averaging each spot’s expression with its adjacent neighbors 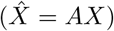 before input to SEPAR. For image-based spatial transcriptomics datasets, which typically contain fewer genes, the raw expression matrices were used directly as input.

SEPAR was implemented with the following default parameters: pattern number *r* = 30, and regularization coefficients *α* = 0.5, *β* = 0.01, and *γ* = 0.5. For downstream analyses, we set the threshold parameters as *ϵ*_*_ = 0.3 for pattern-specific gene detection and *ϵ* = 0.7 for spatially variable gene identification. In spatial domain analysis, the default settings are *N*_1_ = *r* − 5 − *n*_cluster_ and *N*_2_ = 3, where *n*_cluster_ represents the number of clusters. Detailed parameter settings for all datasets analyzed in this study are provided in Supplementary Note 2.

### Competing methods

SEPAR was compared with state-of-the-art methods for different analytical tasks. For spatial domain identification and clustering analysis, we employed STAGATE [13] (https://github.com/zhanglabtools/STAGATE), GraphST [45] (https://github.com/JinmiaoChenLab/GraphST), and BASS [12] (https://github.com/zhengli09/BASS) as competing methods. For spatially variable gene detection, we benchmarked against STAMarker [31] (https://github.com/zhanglabtools/STAMarker), SPARK-X [30] (https://xzhoulab.github.io/SPARK/), and SpatialDE [9] (https://github.com/Teichlab/SpatialDE).

To ensure fair comparison, all data were preprocessed according to the protocols described in the original publications of each competing method. The analyses were performed using the official implementations provided by the original authors, with default parameter settings as specified in their released versions.

## Data availability

The spatial transcriptomics datasets analyzed in this study are publicly available: the human DLPFC dataset can be accessed at spatialLIBD project website (http://research.libd.org/spatialLIBD/); the processed mouse olfactory bulb Stereo-seq data is available at https://doi.org/10.5281/zenodo.8356092; the MERFISH data can be downloaded from https://doi.org/10.5061/dryad.8t8s248; the osmFISH data is available at http://linnarssonlab.org/osmFISH/availability/; the MISAR-seq and spatial CITE-seq datasets are available at https://doi.org/10.5281/zenodo.7480069 and https://www.ncbi.nlm.nih.gov/geo/query/acc.cgi?acc=GSE213264, respectively.

## Code availability

The SEPAR software package is freely available at https://github.com/zerovain/SEPAR under the MIT license. Detailed documentation and tutorials are available at https://SEPAR.readthedocs.io.

## Supplementary information

Supplementary information includes additional data visualization (Supplementary Figs. 1-17), supplementary discussion of SEPAR’s enhanced performance in multi-slice SRT integration (Supplementary Note 1 and Supplementary Fig. 18), detailed parameter settings for different datasets (Supplementary Note 2).

## Acknowledgements

This work has been supported by National Key Research and Development Program of China (No. 2021YFC2701601 to S.Z., No. 2023YFF1204802 to Y.Z.), Science and Technology Commission of Shanghai Municipality (No. 23JC1401000 to S.Z.), National Natural Science Foundation of China (No. 12471350 to S.Z., No. 82071259 to Y.Z.), and National Science and Technology Innovation 2030 Major Program (Grant No. 2021ZD0200100 to Y.Z.).

