## Supplemental figures and notes for "Spatial Metagene Discovery and Associated Molecular Pattern Characterization in Spatial Transcriptomics and Multi-Omics using SEPAR"

### Supplementary Information

April 3, 2025

#### 1 Supplementary Figures

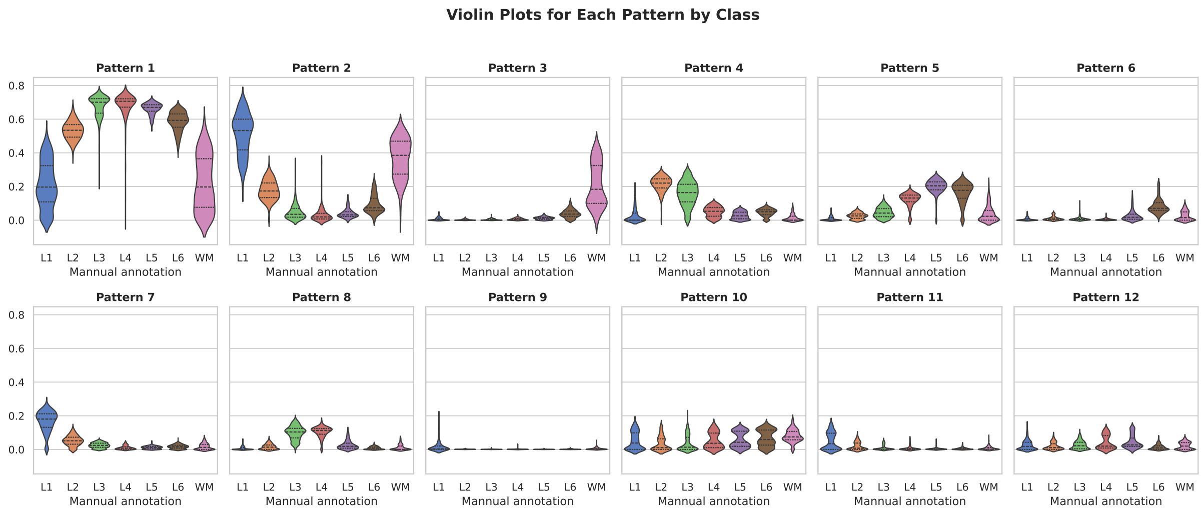


Figure S1: The distribution of expression levels for each pattern illustrated using violin plots across seven manually annotated regions.

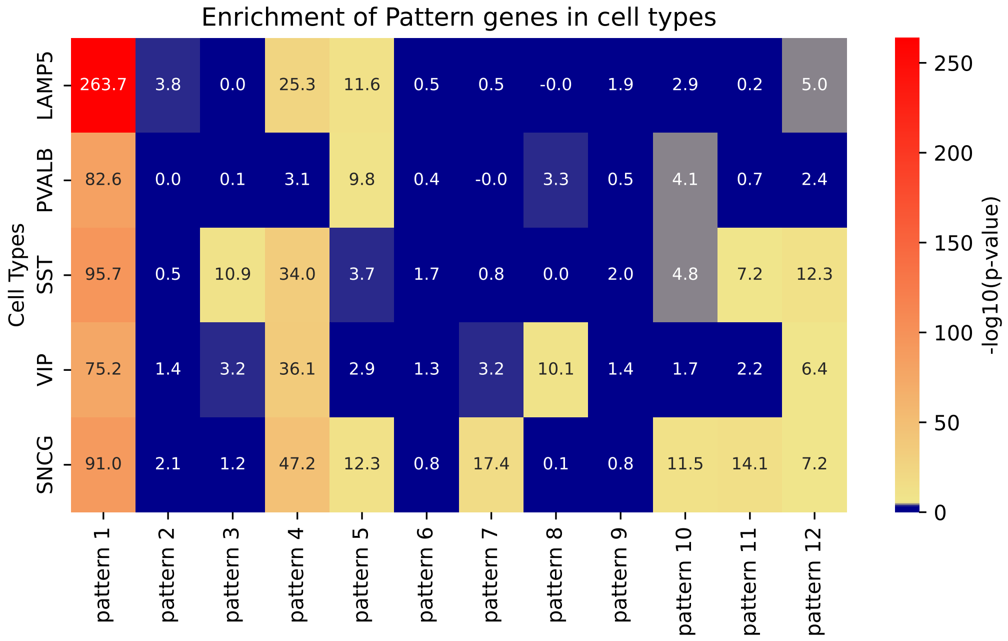


Figure S2: Cell-type enrichment analysis on the gene sets corresponding to each pattern using CellGO, using identifier with InN cell including LAMP5, PVALB, SST, VIP, and SNCG

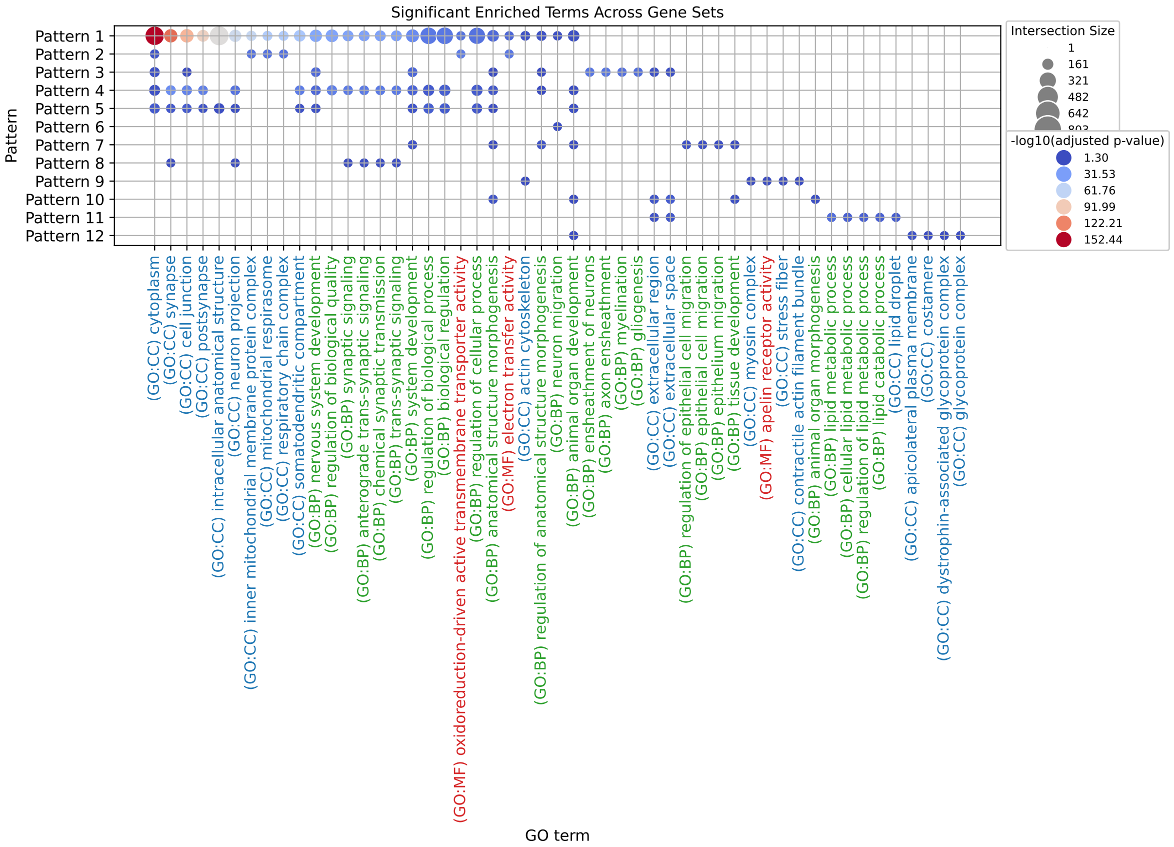


Figure S3: Gene Ontology (GO) term enrichment results for patterns 1-12 identified in the DLPFC dataset, corresponding to 12 specific gene sets. The top five significant GO terms for each set are displayed in bubble charts, with GO: Biological Process, GO: Molecular Function, and GO: Cellular Component colored green, red, and blue, respectively.

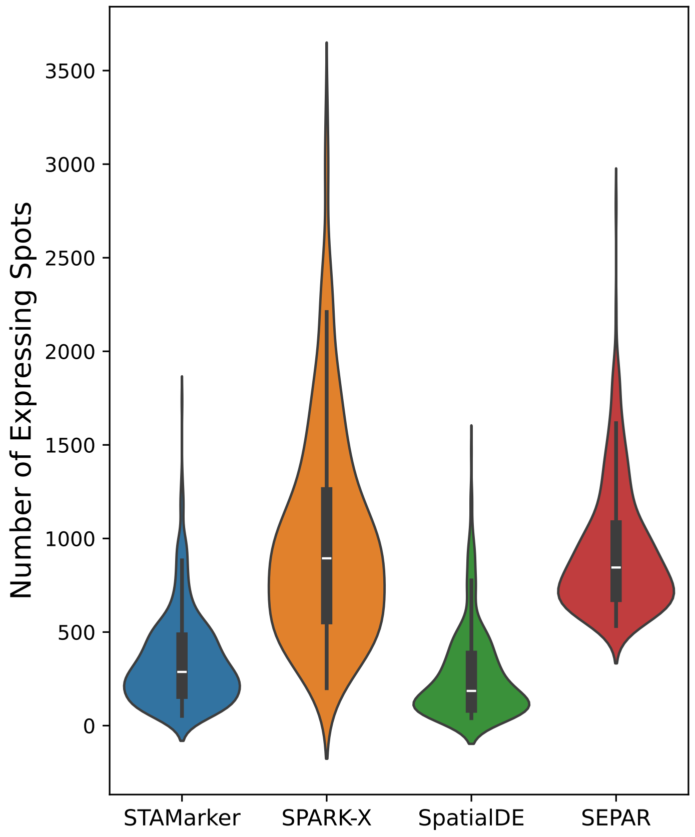


Figure S4: Violin plot of expressing spot number of uniquely identified SVGs for each method on DLPFC dataset.

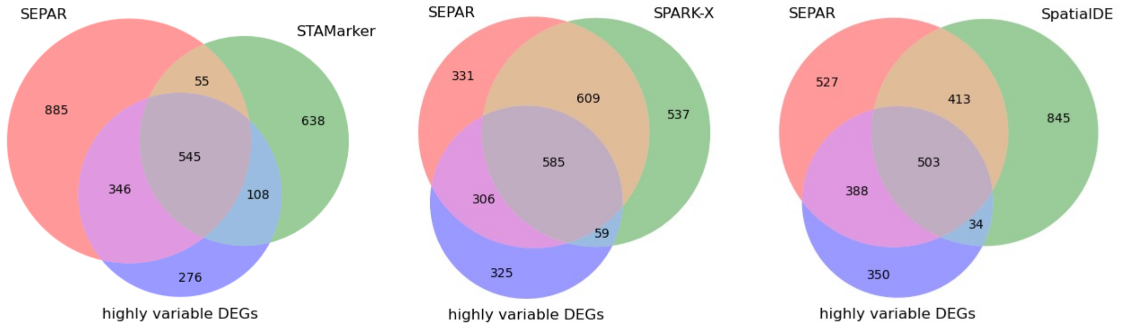


Figure S5: The SEPAR analysis conducted on section 151507 of the DLPFC dataset. A Venn diagram compares SEPAR with other known spatially variable gene (SVG) identification methods such as STAMarker, SPARK-X, and SpatialDE on high variable SVGs.

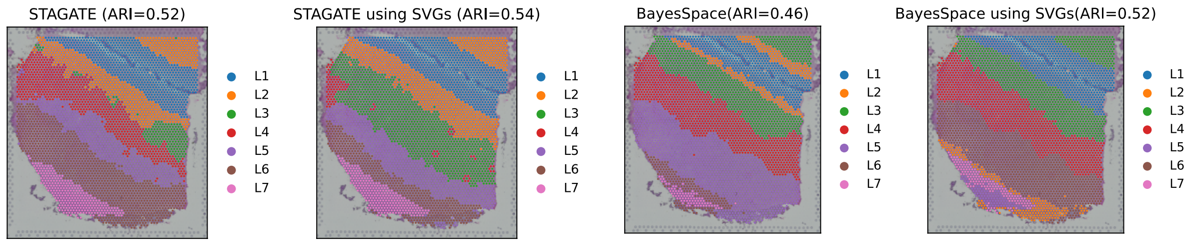


Figure S6: Clustering results of STAGATE and BayesSpace using their default setting(top 3000 HVGs as input for STAGATE and top 2000 HVGs as input for BayesSpace) or using SVGs selected by SEPAR.

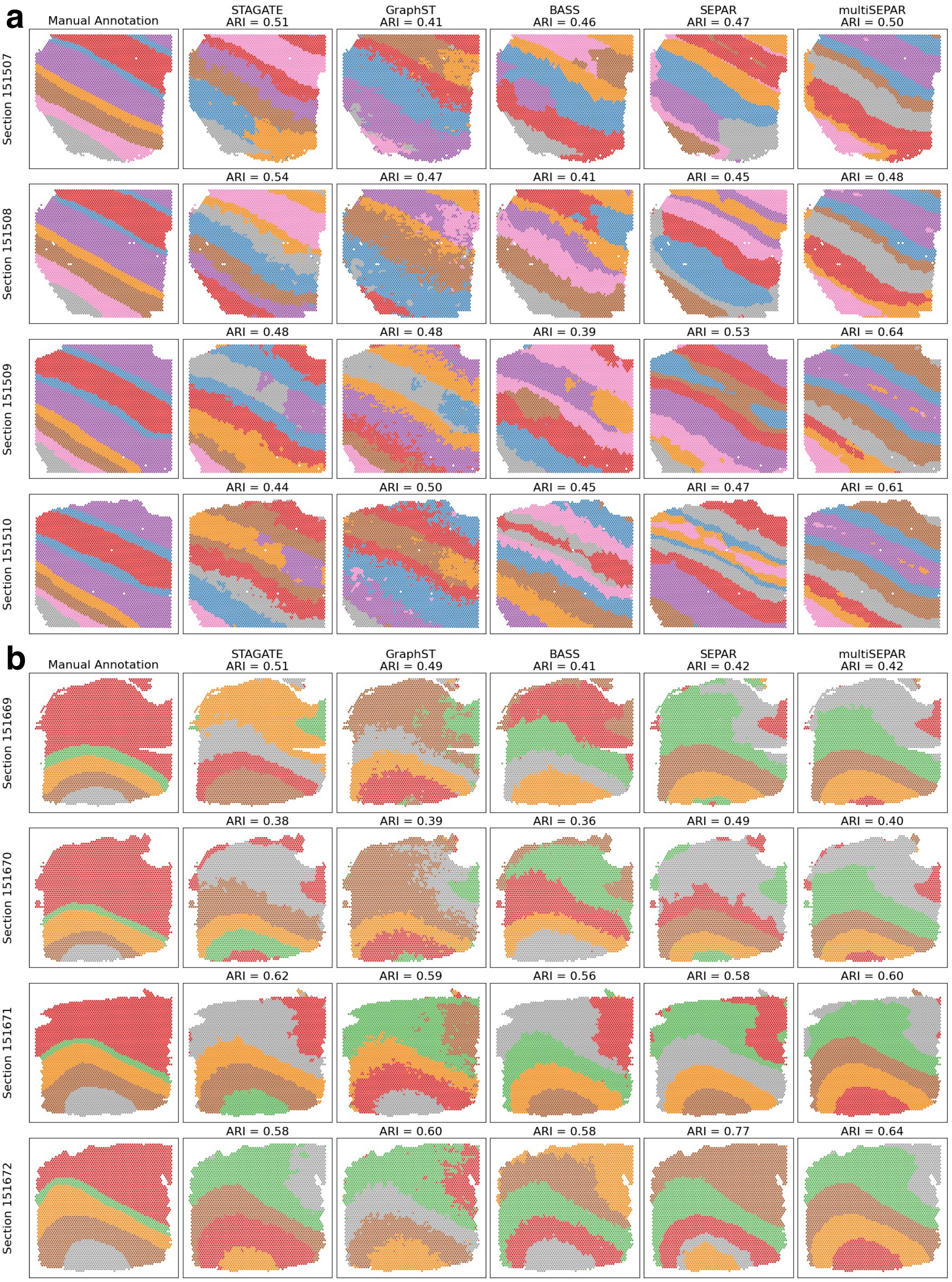


Figure S7: (**a**) The sections 151507-151510 of the DLPFC dataset display manual annotations along with cluster identifications using methods like STAGATE, GraphST, BASS, SEPAR, and SEPARmult. (**b**) Similarly, sections 151669-151672 are analyzed for manual annotations and cluster identifications using the same methods.

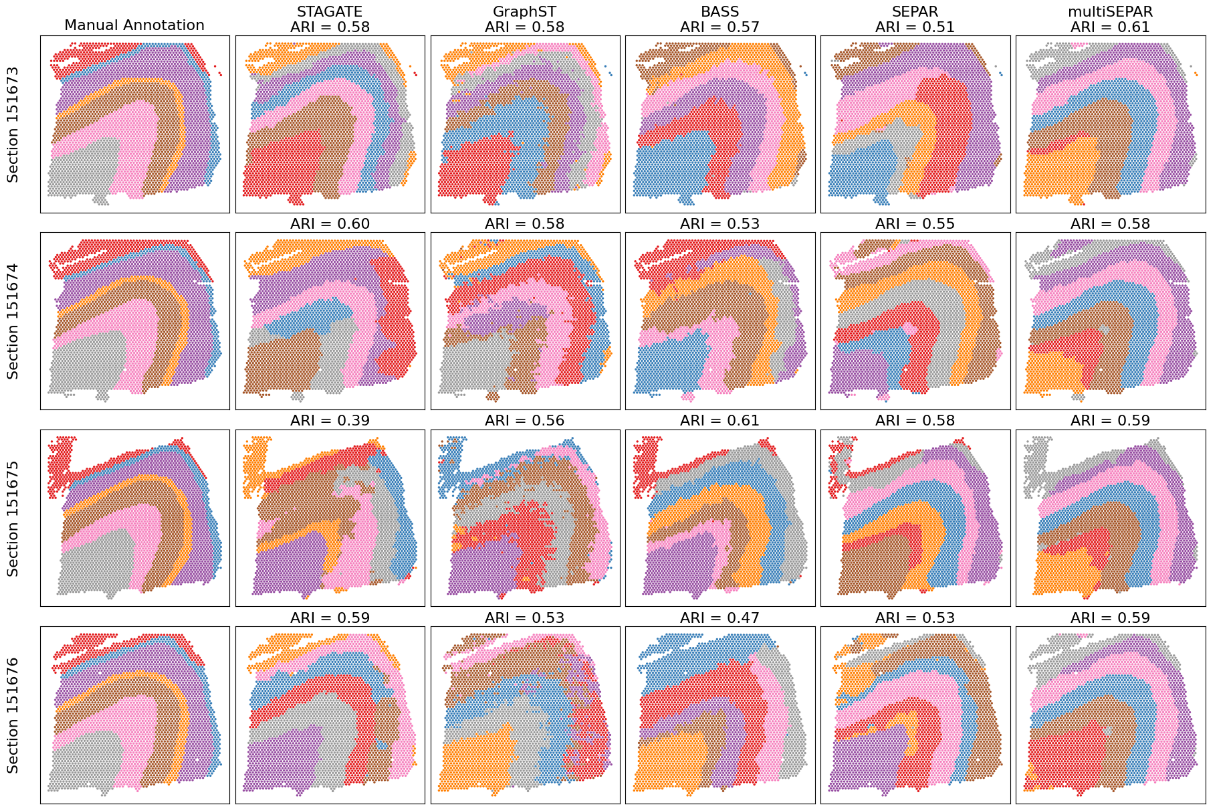


Figure S8: The sections 151673-151676 of the DLPFC dataset display manual annotations along with cluster identifications using methods like STAGATE, GraphST, BASS, SEPAR, and SEPARmult.

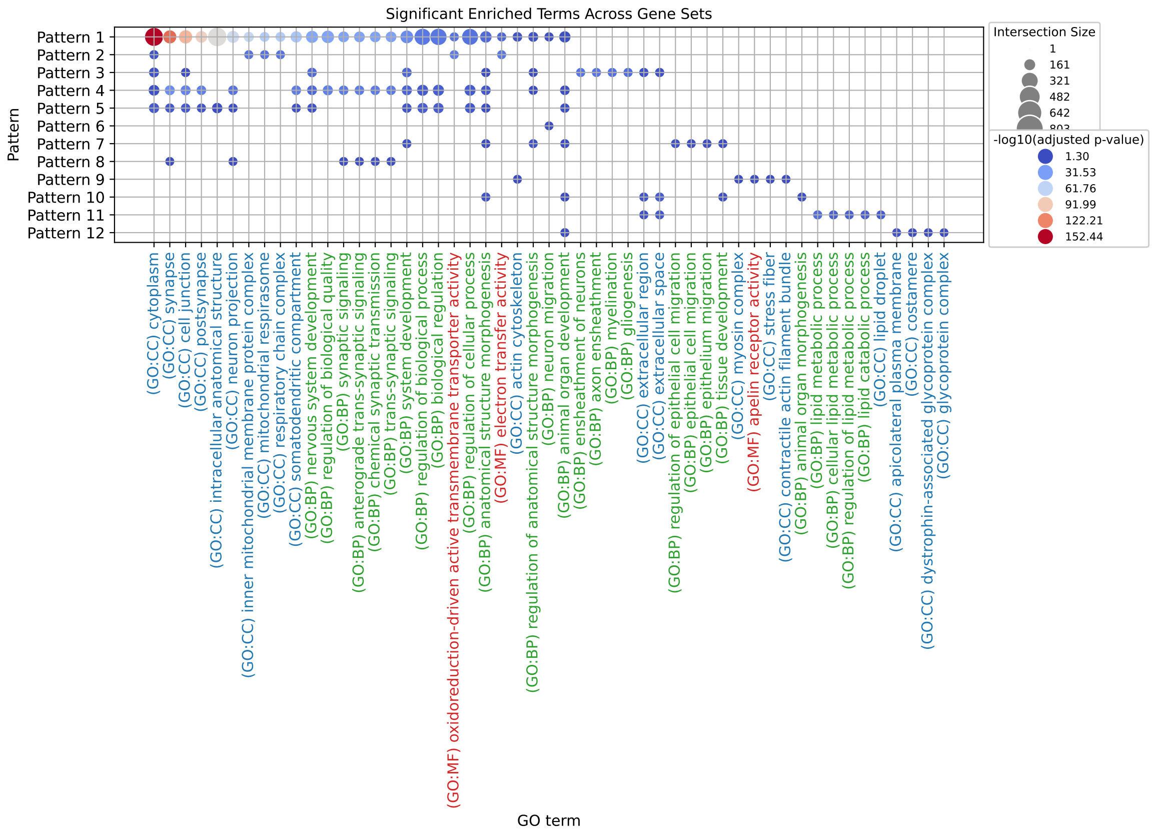


Figure S9: GO term enrichment results are shown for patterns 1-12 identified in the Stereo-seq dataset, with pattern 11 having no significant GO terms due to insufficient gene count. Each set's top five significant GO terms are depicted in bubble charts with color-coding for GO categories.

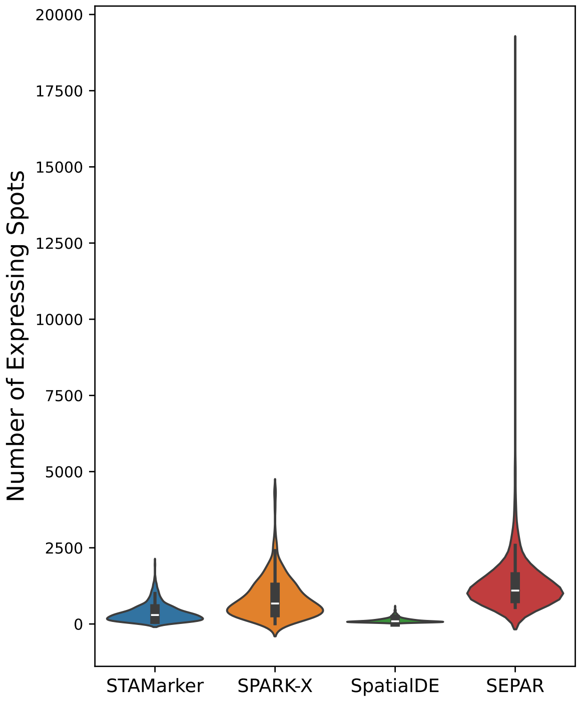


Figure S10: Violin plot of expressing cell number of uniquely identified SVGs for each method on Stereo-seq dataset.

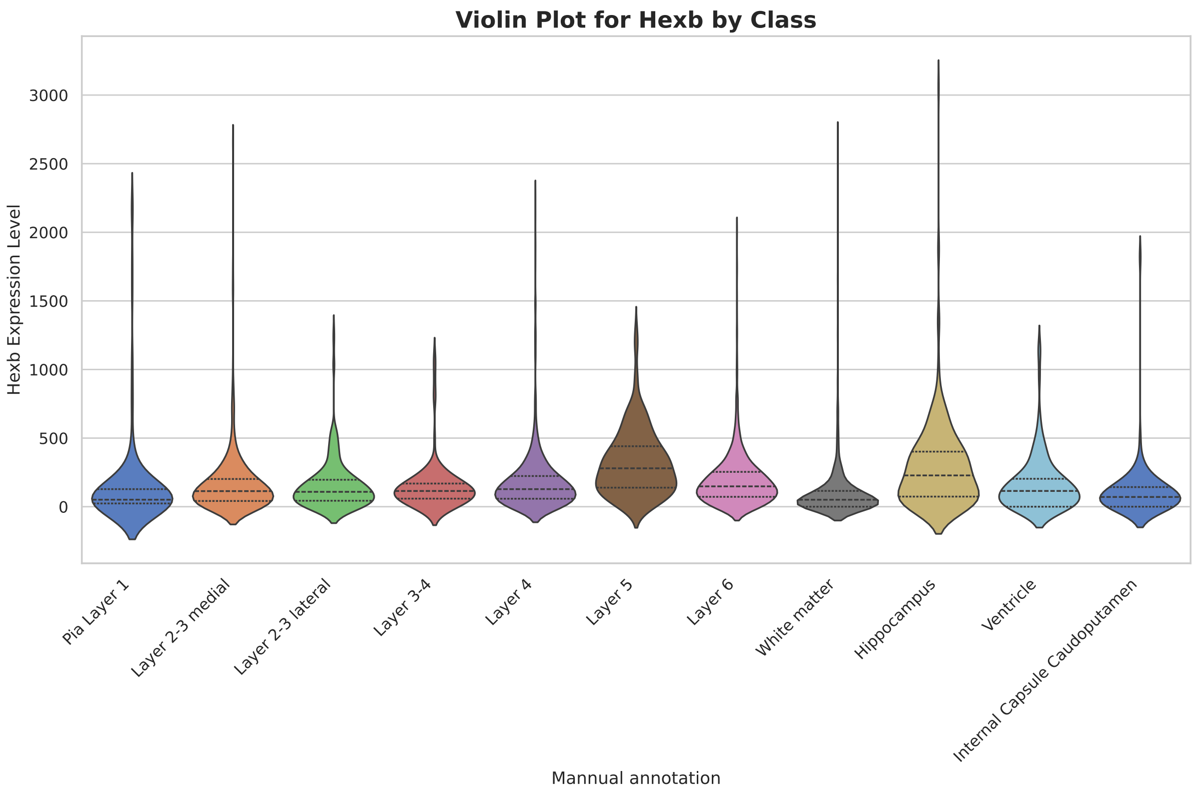


Figure S11: Violin plots compare the expression distribution of the Hexb gene across 11 domains in the osmFISH dataset.

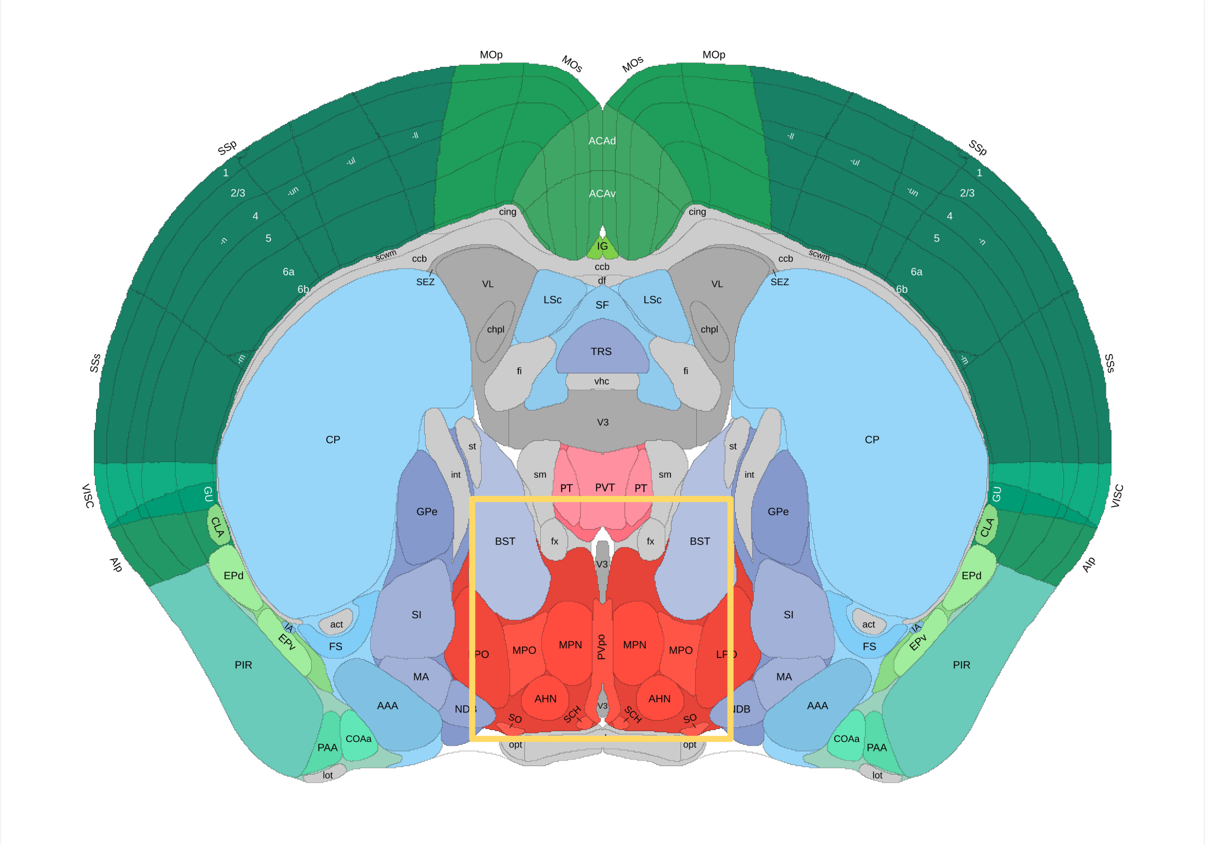


Figure S12: Anatomical reference from the Allen Brain Atlas showing mouse brain structures, with the yellow box delineating the hypothalamic preoptic region that corresponds to the MERFISH-profiled area.

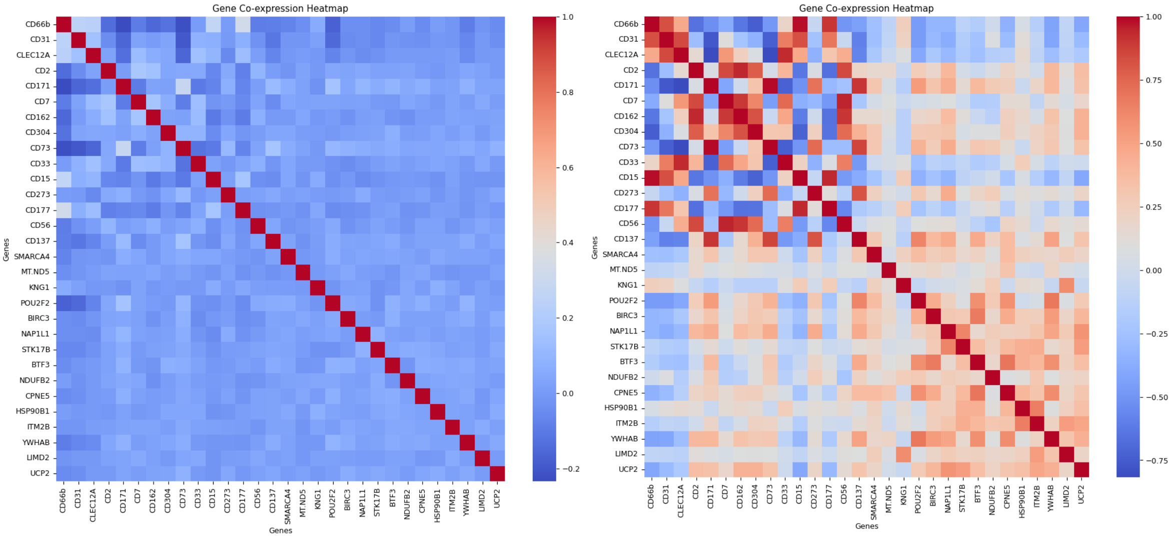


Figure S13: Gene co-expression analysis within the spatial CITE-seq dataset includes: (**a**) Heatmaps representing gene co-expression using raw gene expression data. (**b**) Heatmaps based on spatially refined gene co-expression.

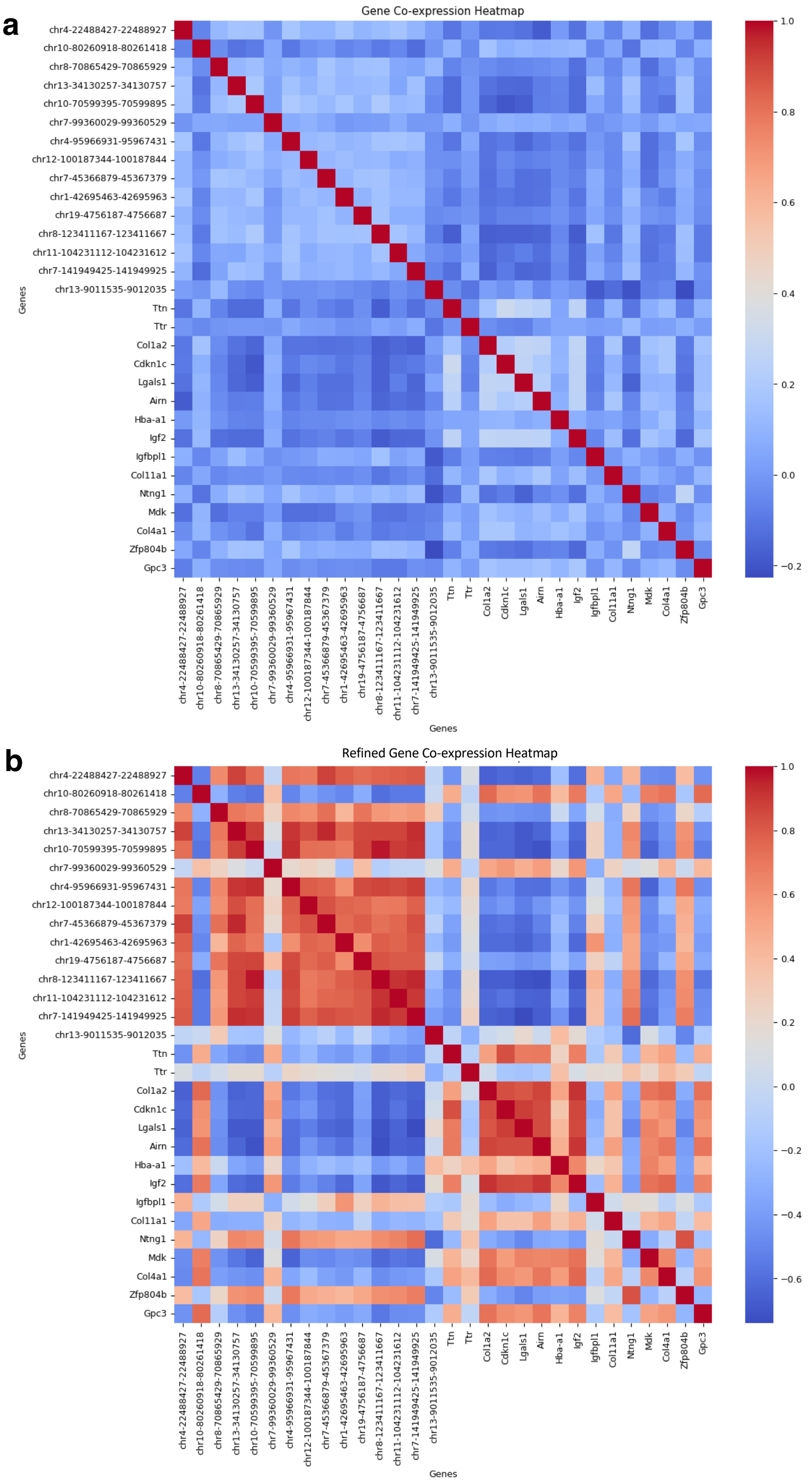


Figure S14: Gene co-expression analysis within the MISAR-seq dataset includes: (**a**) Heatmaps representing gene co-expression using raw gene expression data. (**b**) Heatmaps based on spatially refined gene co-expression.

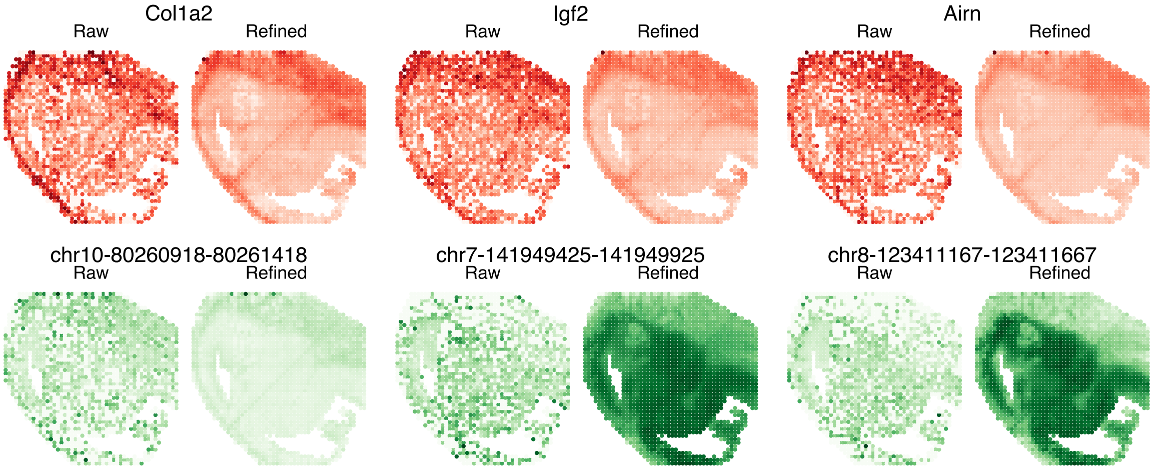


Figure S15: Visualization of three genes and three peaks before and after refinement.

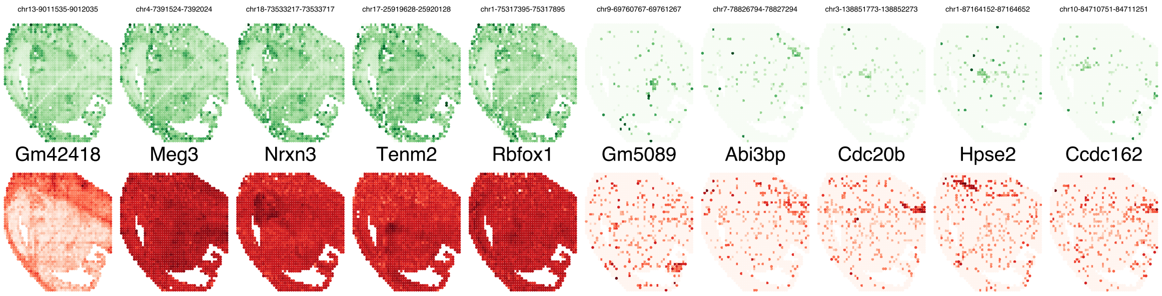


Figure S16: SEPAR's identification of spatial/non-spatial variable peaks and genes in the MISAR-seq dataset.

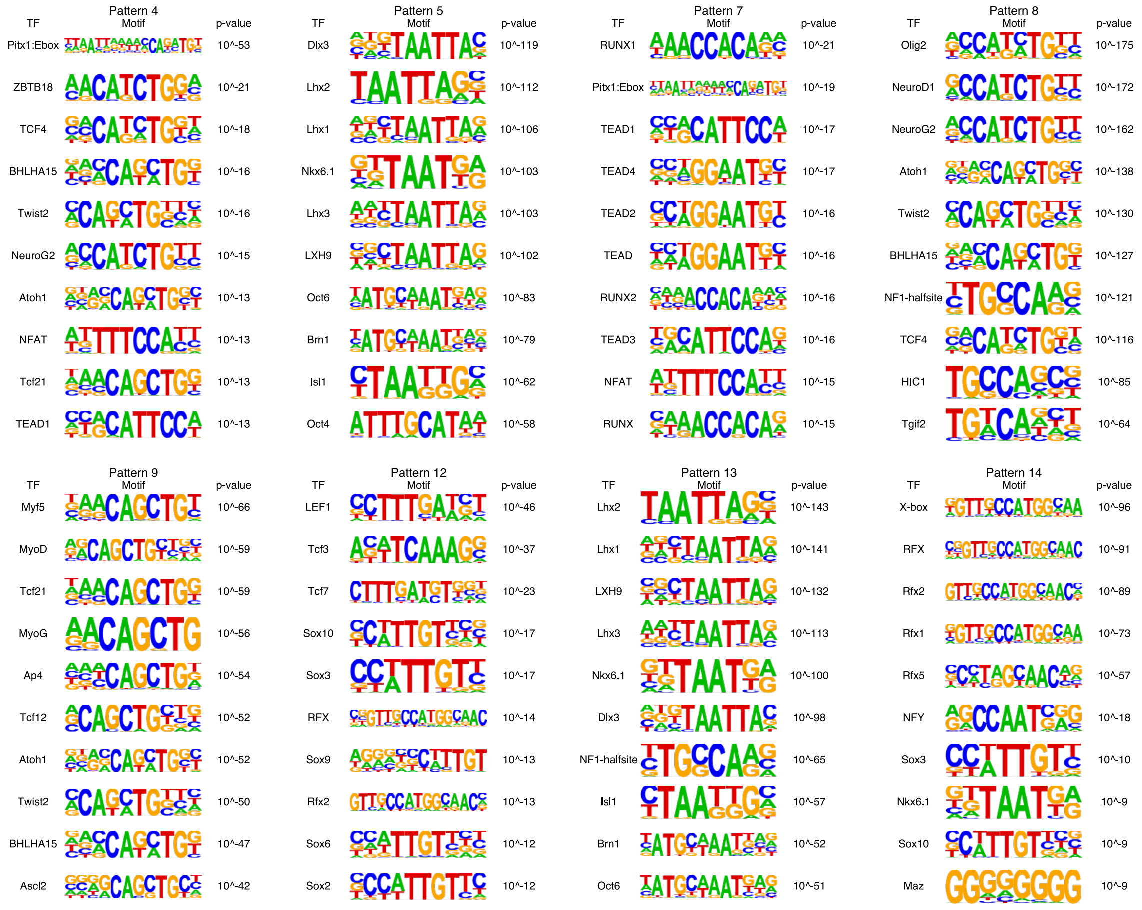


Figure S17: Downstream motif enrichment analysis of MISAR-seq data from E15.5 mouse embryonic brain tissue. HOMER was used to identify enriched transcription factor binding motifs within each pattern-specific peak set identified by SEPAR.

### 2 Supplementary Note 1: Enhanced performance of SEPAR in multislice SRT integration

To demonstrate SEPAR's extensibility to multiple tissue sections, we developed SEPARmult for multislice SRT integration analyses. SEPARmult extends the original SEPAR framework to simultaneously analyze multiple slices of tissue SRT data from the same sample source ($X_s, s = 1, \ldots, M$). For each slice, we compute a slice-specific Laplacian matrix $L_s$ and formulate the optimization problem as:

$$\min_{W_s \geq 0, H \geq 0} \sum_{s=1}^{M} \left( \|X_s - W_s H\|_2^2 + \alpha \operatorname{tr}(\operatorname{diag}(PSS) W_s^T L_s W_s) + \beta \|W_s\|_1 \right) + \frac{\gamma}{2} \sum_{i<j} (\langle h_i, h_j \rangle)^2. \tag{1}$$

The optimization algorithm follows similar principles as the original SEPAR method (**Methods**). This formulation allows SEPARmult to leverage information across multiple tissue sections while maintaining slice-specific spatial relationships.

We evaluated SEPARmult using the DLPFC dataset, which comprises 12 tissue slices from human dorsolateral prefrontal cortex [1]. When analyzing spatial domain identification across these samples, SEPAR achieved a median ARI of 0.520 in single-slice mode. SEPARmult significantly improved this performance to a median ARI of 0.590, outperforming other state-of-the-art methods including STAGATE (median ARI = 0.525), GraphST (median ARI = 0.515), and BASS (median ARI = 0.465) (Fig. S18a).

To illustrate the improvement in spatial domain identification, we examined four adjacent samples (sections 151673-151676). The results demonstrate that SEPARmult (Fig. S18c) achieved more consistent and biologically meaningful domain identification compared to single-slice analysis using SEPAR (Fig. S18b). This enhancement in performance can be attributed to SEPARmult's ability to leverage cross-section information, resulting in more robust and stable spatial domain identification.

These results highlight the advantage of integrating information across multiple tissue sections, particularly for studies involving serial sections or replicate samples. The improved performance of SEPARmult suggests its potential utility in large-scale spatial transcriptomics studies where multiple tissue sections need to be analyzed collectively.

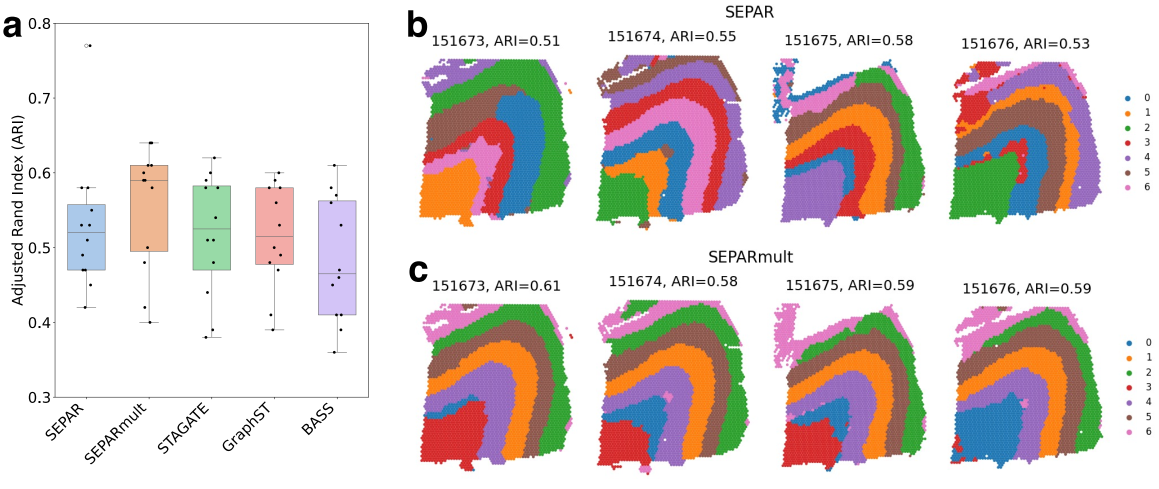


Figure S18: **Performance evaluation of SEPARmult on DLPFC dataset.** (**a**) Comparison of clustering performance across methods. Boxplots show the distribution of ARI values for spatial clustering results using SEPAR, SEPARmult, STAGATE, GraphST, and BASS across 12 tissue sections. (**b**) Spatial clustering results using SEPAR on individual sections 151673-151676. (**c**) Spatial clustering results using SEPARmult on sections 151673-151676, demonstrating improved consistency and stability in domain identification.

### 3 Supplementary Note 2: Parameter settings for different datasets

Table 1: Hyperparameter settings for SEPAR across different spatial datasets and experimental conditions

| Dataset | Slice | $r$ | $\alpha$ | $\beta$ | $\gamma$ | $N_1$ | $N_2$ |
|---|---|---|---|---|---|---|---|
| | 151507 | 30 | 0.7 | 0.01 | 0.3 | 16 | 5 |
| | 151508 | 30 | 0.7 | 0.01 | 0.3 | 16 | 5 |
| | 151509 | 30 | 0.7 | 0.01 | 0.3 | 16 | 5 |
| | 151510 | 30 | 0.7 | 0.01 | 0.3 | 16 | 5 |
| | 151669 | 30 | 0.3 | 0.02 | 0.3 | 18 | 5 |
| | 151670 | 30 | 0.3 | 0.02 | 0.3 | 18 | 5 |
| DLPFC | 151671 | 30 | 0.3 | 0.02 | 0.3 | 18 | 5 |
| | 151672 | 30 | 0.3 | 0.02 | 0.3 | 18 | 5 |
| | 151673 | 30 | 0.3 | 0.02 | 0.3 | 16 | 5 |
| | 151674 | 30 | 0.3 | 0.02 | 0.3 | 16 | 5 |
| | 151675 | 30 | 0.3 | 0.02 | 0.3 | 16 | 5 |
| | 151676 | 30 | 0.3 | 0.02 | 0.3 | 16 | 5 |
| | multi-slice | 30 | 0.5 | 0.003 | 0.5 | 13 | 5 |
| Stereo-seq | – | 30 | 0.8 | 0.05 | 0.5 | 16 | 4 |
| osmFISH | – | 30 | 1.0 | 0.05 | 0.01 | 0 | 0 |
| MERFISH | – | 30 | 1.0 | 0.05 | 0.01 | 5 | 7 |
| CITE-seq | – | 30 | 0.5 | 0.01 | 0.5 | 18 | 7 |
| MISAR-seq | – | 30 | 0.5 | 0.01 | 0.5 | 15 | 1 |
